# Internal Neural Representations in Task Switching Guided by Context Biases

**DOI:** 10.1101/2023.07.24.550365

**Authors:** Santiago Galella, Salva Ardid

## Abstract

Our brain can filter and integrate external information with internal representations to accomplish goal-directed behavior. The ability to switch between tasks effectively in response to context and external stimuli is a hallmark of cognitive control. Task switching occurs rapidly and efficiently, allowing us to perform multiple tasks with ease. Similarly, artificial intelligence can be tailored to exhibit multitask capabilities and achieve high performance across domains. In this study, we delve into neural representations learned by *task-switching* feedforward networks, which use task-specific biases for multitasking mediated by *context inputs*. Task-specific biases are learned by alternating the tasks the neural network learns during training. By using two-alternative choice tasks, we find that *task-switching* networks produce representations that resemble other multitasking paradigms, namely *parallel* networks in the early stages of processing and *independent* subnetworks in later stages. This transition in information processing is akin to that in the cortex. We then analyze the impact of inserting task contexts in different stages of processing, and the role of its location in the alignment between the task and the stimulus features. To confirm the generality of results, we display neural representations during task switching for different task and data sets. In summary, the use of *context inputs* improves the interpretability of feedforward neural networks for multitasking, setting the basis for studying architectures and tasks of higher complexity, including biological microcircuits in the brain carrying out context-dependent decision making.

## Introduction

Action selection is the result of interactions between different aspects, such as external inputs, specific contexts and goals (Monsell, 2003). Alternating tasks efficiently, *task switching*, is a hallmark of cognitive control. In the brain, the main area that has been related with task switching and cognitive control is the prefrontal cortex (PFC) (K. Johnston et al., 2007). The PFC is believed to contain different microcircuits that provide rule encoding and top-down regulation (Menon & D’Esposito, 2022) to other brain areas transitioning from sensory processing to action selection. However, the exact mechanisms are not fully understood. One hypothesis, the gating theory (Miller & Cohen, 2001), posits that the PFC mediates the activation of different neural pathways by sending top-down context modulations. Many neural circuit models have been proposed to explain task switching in the brain (J. D. Cohen et al., 1990; Botvinick et al., 2001; Rougier & O’Reilly, 2002; Ardid & Wang, 2013). More recently, artificial neural networks have been analyzed as models of neural computation,

Internal Neural Representations in Task Switching Guided by Context Biases many findings suggesting their practicality for assessing brain theories (Richards et al., 2019), including the study of context in cognitive control in humans and machines (Mante et al., 2013; Yang et al., 2019).

Two main paradigms have been compared in terms of information processing in artificial neural networks (Ruder, 2017; Crawshaw, 2020): According to *independent* processing, different modules or subnetworks are specialized in isolation one from the others, ensuring the allocation of specific resources for distinct modalities and avoiding interference. However, *independent* processing relies on duplicating resources to execute multiple tasks in isolation. In *parallel* processing, in contrast, distinct modalities are processed simultaneously within a same network. Through the course of learning, *parallel* networks can benefit from shared internal representations between tasks (Caruana, 1997). Shared representations help regularizing and promoting generalization, improving network performance across domains (Ruder, 2017). However, *parallel* networks strictly require executing all tasks at once, which can be consuming and inefficient.

Inspired by the role of the PFC in goal-directed behavior, we introduce the *task-switching* network, which aligns with task-specific context-dependent decision making. During training, *task-switching* networks learn specific context biases for each task. Yet, abundant shared representations remain in *task-switching* networks due to shared weight parameters across tasks. Context biases, nevertheless, help routing subnetworks in downstream layers that execute either task, regulating the mode at which the network operates: *parallel* or *independent*.

Previous research explored using task contexts in neural networks to analyze multitask efficiency (Musslick et al., 2017; Ravi et al., 2020), to compare neural geometry with that in the brain (Flesch, Juechems, et al., 2022), or as task embeddings for multitasking in computer vision applications (Sun et al., 2021). In this study, however, we focus on comparing context-dependent decision making with *independent* and *parallel* processing by analyzing internal neural representations in *task-switching* networks. For this, we apply population analysis techniques developed to investigate how neural computations associate stimulus features with task demands (Kriegeskorte et al., 2008; Jazayeri & Ostojic, 2021), and to investigate internal neural representations in artificial neural networks (Bengio et al., 2013; Ito & Murray, 2021; Ito et al., 2022; Flesch, Juechems, et al., 2022).

First, we compare the representations learned by *independent, parallel* and *task-switching* networks (Section *Task-switching* networks operate in a midway regime between *independent* and *parallel* networks). We observe that *task-switching* networks produce representations that resemble *parallel* networks in the first layers of processing, whereas representations transition toward those of *independent* subnetworks in latter layers. Remarkably, we show that *task-switching* networks, by using context biases, potentiate shared internal representations further than *parallel* networks when learning several tasks simultaneously. We also show that *independent* and *task-switching* in contrast to *parallel* networks underlie neural representations that disentangle the reconstruction of the task, increasing the interpretability of the model. Unexpectedly, *task-switching* networks show that there is a continuity in information processing, with *independent* and *parallel* networks being the two opposite extremes. In fact, *task-switching* networks transition from one to the other based on context allocation (Section Context location determines the transition between *independent* and *parallel* processing).

We next analyze the impact of the context positioning and of its magnitude at different stages of neural processing. Our results show that contexts introduced at mid processing drive neural activity more strongly, akin to the strength of attentional modulations in sensory cortices (Section Context impact is maximal at intermediate layers). Intermediate layers in our model still show *parallel* processing, summarized by task similarity (Section *Task-switching* networks encode task similarity), but initiate the transition towards specific task execution. After that, we show that trained *task-switching* networks are able to extrapolate, generalizing task biases for a wide range of unseen context strengths, while the network still displays robust behavior (Section *Task-switching* networks are robust against *physiological* context modulations). Finally, we employ clustering techniques to identify neural ensembles in *task-switching* networks according to their selectivity to sensory features and task information (Section Neural selectivity emerges to resolve task execution).

The results in this article demonstrate that the use of *context inputs* significantly enhances the interpretability of feedforward neural networks supporting multitasking. Particularly relevant is the theoretical approach introduced in methods (see Section Final representation motifs), based on generalizing the definition of congruency (see Section Generalized congruency), that is able to predict with high accuracy the final internal representations of *individual, parallel* and *task-switching* networks. We hope that the approach and insights presented in this manuscript serve to better understand multitasking in network architectures of higher complexity, including neural circuits in the brain materializing context-dependent decision making.

## Results

Neural representations in brain circuits refer to patterns of activity associated with information processing, for instance encoding features of the environment (Vilarroya, 2017). Similarly, in neural network models, we can use neural representations to refer to different patterns of activity arising from internal processing, for instance for different stimulus inputs. We can investigate the representations learned by neural networks seeking direct mappings between stimuli and their representations, but also by examining the similarities in representations elicited by different stimuli (Edelman, 1998).

### *Task-switching* networks operate in a midway regime between *independent* and *parallel* networks

In systems neuroscience, a widely used methodology to estimate the similarity of different stimuli is the representational similarity analysis (RSA) (Kriegeskorte et al., 2008). In this study, we delve into the similarity between neural representations of *independent, parallel* and *task-switching* networks (Fig 1). Detailed information of their architecture is described in the Section Neural network architectures.

**Figure 1:**
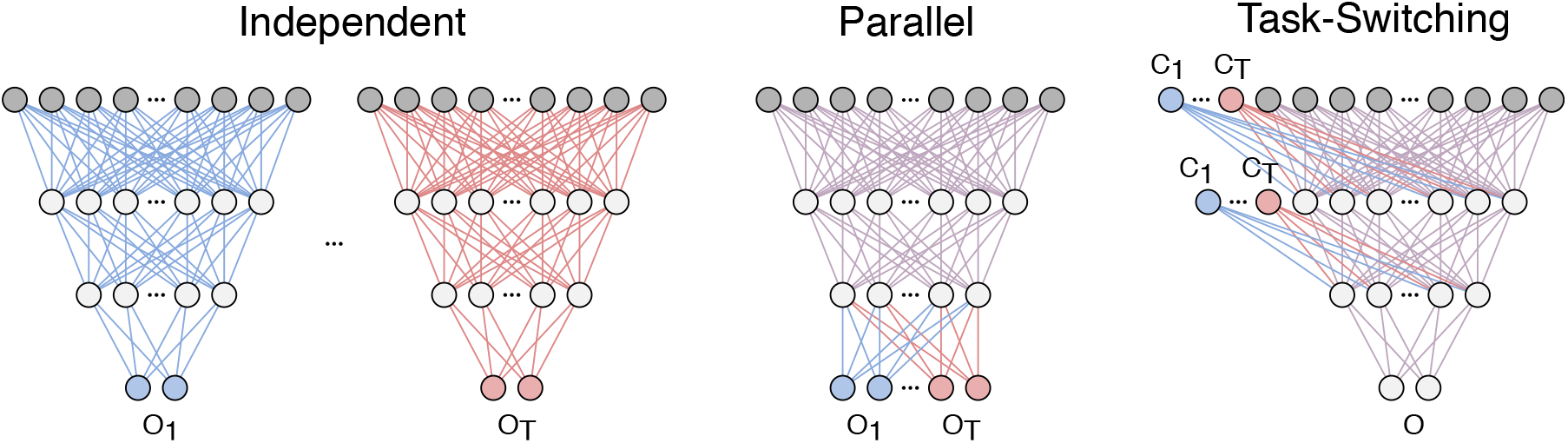
Architectures for different types of information processing. *O*_1_ - *O*_*T*_ represent the outputs for each task. For *independent* processing, tasks are processed separately by isolated modules, whereas for *parallel* processing the same module processes all tasks simultaneously. For the *task-switching* network, the task changes according to which context input from *C*_1_ - *C*_*T*_ is activated.

To study internal representations in these neural network models, we used the two-alternative choice with two tasks, *parity* and *value* (see Sections Two-alternative choice tasks and Training for details). We first visualized the representational dissimilarity matrices (RDMs) for the different architectures (Fig 2a). The RDM describes the (dis-)similarity between activation patterns. Using this technique, we can contrast activation patterns in different conditions, such as patterns for different inputs in the same task, same inputs in distinct tasks, and different inputs in distinct tasks.

**Figure 2:**
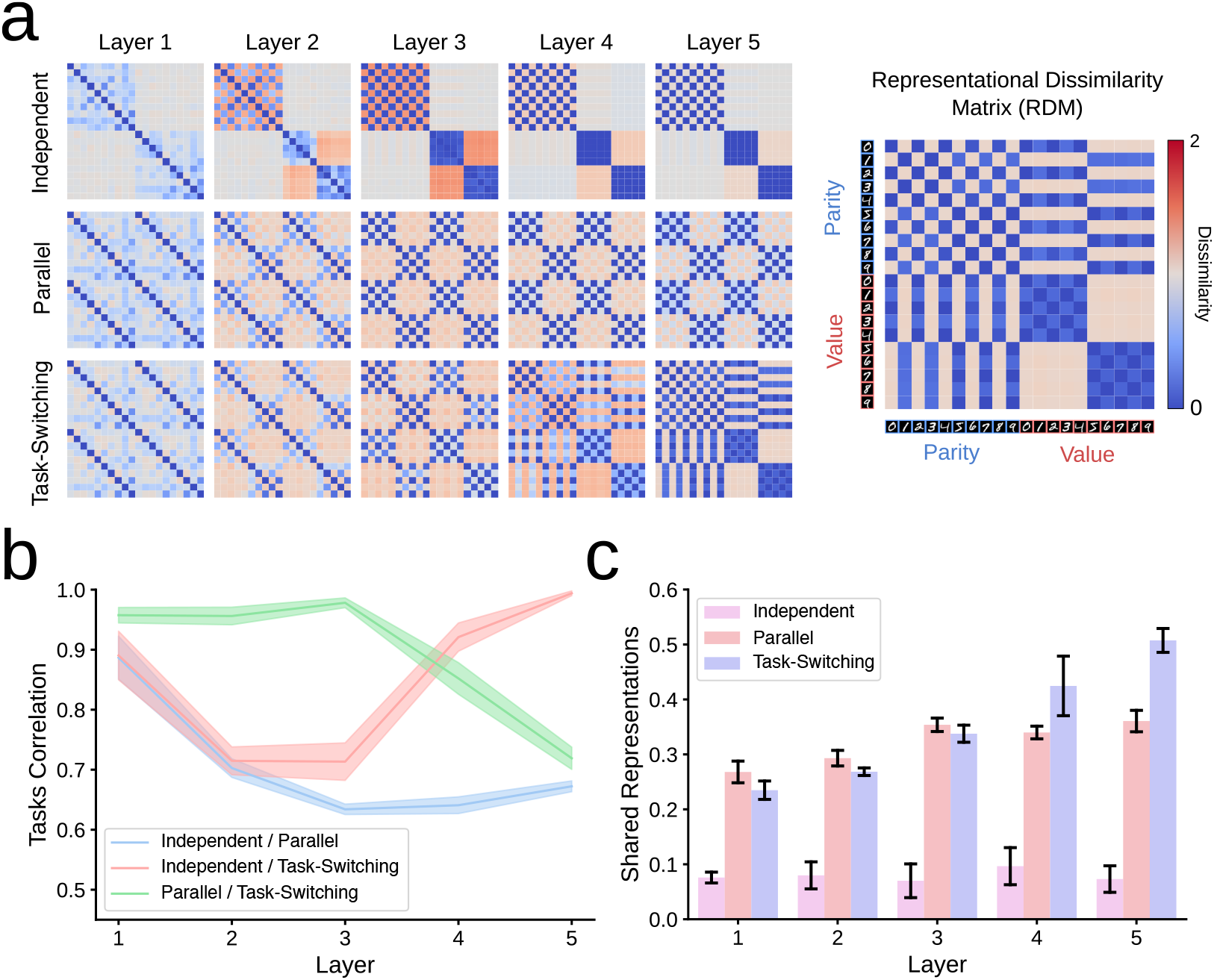
Neural representations for *independent, parallel* and *task-switching* networks for two distinct two-alternative choice tasks based on digit processing: {*parity, value*}. (a) Average representational dissimilarity matrix (RDM) for the three architectures across 10 different initializations (*runs*). (b) Task correlation between models measured by the Pearson correlation between RDMs’ main diagonal blocks (Mean and SD over 10 different runs). (c) Shared representations between different tasks in each model measured by the sum of absolute correlation values in RDMs’ off-diagonal blocks (Mean and SD over 10 runs).

The RDM was built by grouping, in terms of stimulus inputs (e.g., digit images), the neural activations of each layer. Through all tasks and for each distinct stimulus input (e.g., distinct digit label in 0–9), we averaged the neural activity and computed their dissimilarity (see Section Representational dissimilarity matrix for further details). The final RDM is a symmetric matrix where the main-diagonal blocks show the dissimilarity in red (similarity in blue) of activations within each task (main diagonal blocks) and between tasks (off-diagonal blocks).

By inspecting the resulting RDMs (Fig 2a), task-specific responses can be directly decoded in the final stage of internal processing (layer 5) for *independent* and *task-switching* networks. However, this is not the case for *parallel* networks. Intriguingly, the emerging pattern in *parallel* networks is also noticed in *task-switching* networks up to intermediate processing (layer 3). However, the patterns thereafter evolve differently between the two networks.

From RDMs, we obtained the RSA that provides an overall degree of similarity between the representations of the distinct networks. The RSA was estimated analyzing the pairwise Pearson correlation between RDM’s main diagonal blocks (Fig 2b). We computed the mean Pearson correlation and its SD across different training initializations (*n* = 10 *runs*). These results confirm that *task-switching* networks behave very similarly to *parallel* networks in the first stages of processing, yet *task-switching* networks then evolve differently to end up behaving akin to *independent* networks. To better understand these patterns see the theoretical approach we developed in Section Final representation motifs.

Next, we developed a metric to estimate shared representation from the similarity of representations across tasks (Caruana, 1997), which we applied to each model and layer (Fig 2c). Shared representations were derived from RDM’s off-diagonal blocks (see Section Shared representations). Results show that *parallel* and *task-switching* networks show significant shared representations, indicative of networks learning common representations for the different tasks. In contrast, the results for the *independent* network only show noise correlations since the different subnetworks composing *independent* processing are mutually disconnected.

To test the generality of these results, we applied an identical approach to (i) a different dataset: the two-alternative choice tasks based on the processing of letters (S1 Fig S1, (ii) an increased number of tasks: five distinct two-alternative choice tasks based on digit processing (S1 Fig S2). Results were robust in all cases and conditions: *task-switching* networks resemble *parallel* and *independent* in early and advanced processing, respectively.

Remarkably, shared representations diminished after increasing the number of distinct two-alternative choice tasks (Fig 2c vs S1 Fig S2b). Yet, the *task-switching* network, unlike the *parallel* network, boosted shared representations in the final processing stage. In this analysis, we specifically inspected the representations learned for *parity* and *value*, when the network was trained for two tasks, and compared them with those after training five tasks: {*parity, value*} plus {*prime, fibonacci, multiples of 3*}. In the latter, we cropped the RDM for *parity* and *value* to ease the comparison (S1 Fig S3). Even though intermediate processing was treated differently in the two conditions, the final representation motif did not vary between the two conditions (S1 Fig S3, layer 5).

Section Final representation motifs relates the final stage of processing patterns in the RDM with the generalized congruency between tasks and stimuli (see Section Generalized congruency). This theoretical approach also specifies a theoretical convergence value of shared representations. The results from this analysis shows that the same underlying principles shape internal neural representations in *task-switching* and *parallel* networks regardless of the number of tasks.

The discrepancy in task scalability between task-switching and *parallel* networks (Fig 2c vs S1 Fig S2b) relies on the fact that *parallel* networks cannot resolve the task until reaching the output layer, when independent synaptic connections map neural activity to different task responses *O*_1_ - *O*_*T*_ (see Fig 1). Training minimizes the amount of errors, by shaping the activations to encode the average task similarity. This encoding becomes progressively less efficient as the number of tasks increases, which is a limitation of *parallel* networks (see S1 Fig S20). For a deeper understanding of shared representations in *parallel* and *task-switching* networks see Section Final representation motifs).

As a control, we assessed the performance of the three architectures by analyzing the accuracy of the tasks. The major intention of this analysis was to show that the accuracy in all networks was robust: high and similar accuracy, regardless of the architecture (S1 Fig S4a). Only when performance was evaluated simultaneously for the two tasks (see Section Two-alternative choice tasks), minor differences reached significance between *independent* and *parallel* networks, and between *independent* and *task-switching* networks (Wilcoxon Signed-

Rank Test, *p <* 0.001). This might be explained in terms of parameter sharing, i.e., weights and biases that are shared between tasks in *parallel* and *task-switching* networks, which may regularize the network performance (Ruder, 2017). However, this represents a rather small size effect. We conducted the same analysis when using five tasks (S1 Fig S4b), as well as applied to two tasks using letters instead of digits (S1 Fig S5), with results in all conditions showing a similar trend.

### Context location determines the transition between *independent* and *parallel* processing

Research suggests that the brain benefits from *parallel* processing underlying sensory processing (Rousselet et al., 2002), but goal-directed behavior, on the other hand, relies on mechanisms controlling covert selective attention and overt action selection. These mechanisms are better captured by task-specific context-dependent decision-making, possibly due to cognitive and physical limitations in the number of actions we can perform in *parallel* (Sigman & Dehaene, 2008). When performing many tasks, the brain can swift between *parallel* vs. *independent* processing, depending of task conditions (Fischer & Plessow, 2015). Here, we investigated the impact of context inputs in determining how *task-switching* networks transition between *parallel* and *independent* processing. To gain higher precision in the evolution of neural representations at different processing stages, we increased the size of the models from 5 to 10 layers (See Section Training).

We compared *independent* and *parallel* networks with *task-switching* networks, in which contexts were added in the first, in the last, or in every hidden layer of the network. For simplicity, we refer to these networks as *First, Last* and *All* layers, respectively. We analyzed task correlations between models across layers to quantify the impact of context inputs with respect to their location.

First, we compared how the context position in the (*First* and *Last*) layers affects the representations compared to the *All* layers condition (Fig 3a). At early processing stages, *Last* and *All* layers locations behave very much alike, but the pattern is reversed later during advanced processing. From this analysis, we infer that contexts in the first layers are not playing an important role in the *All* layers condition since the similarity with the *Last* layer condition is very high.

**Figure 3:**
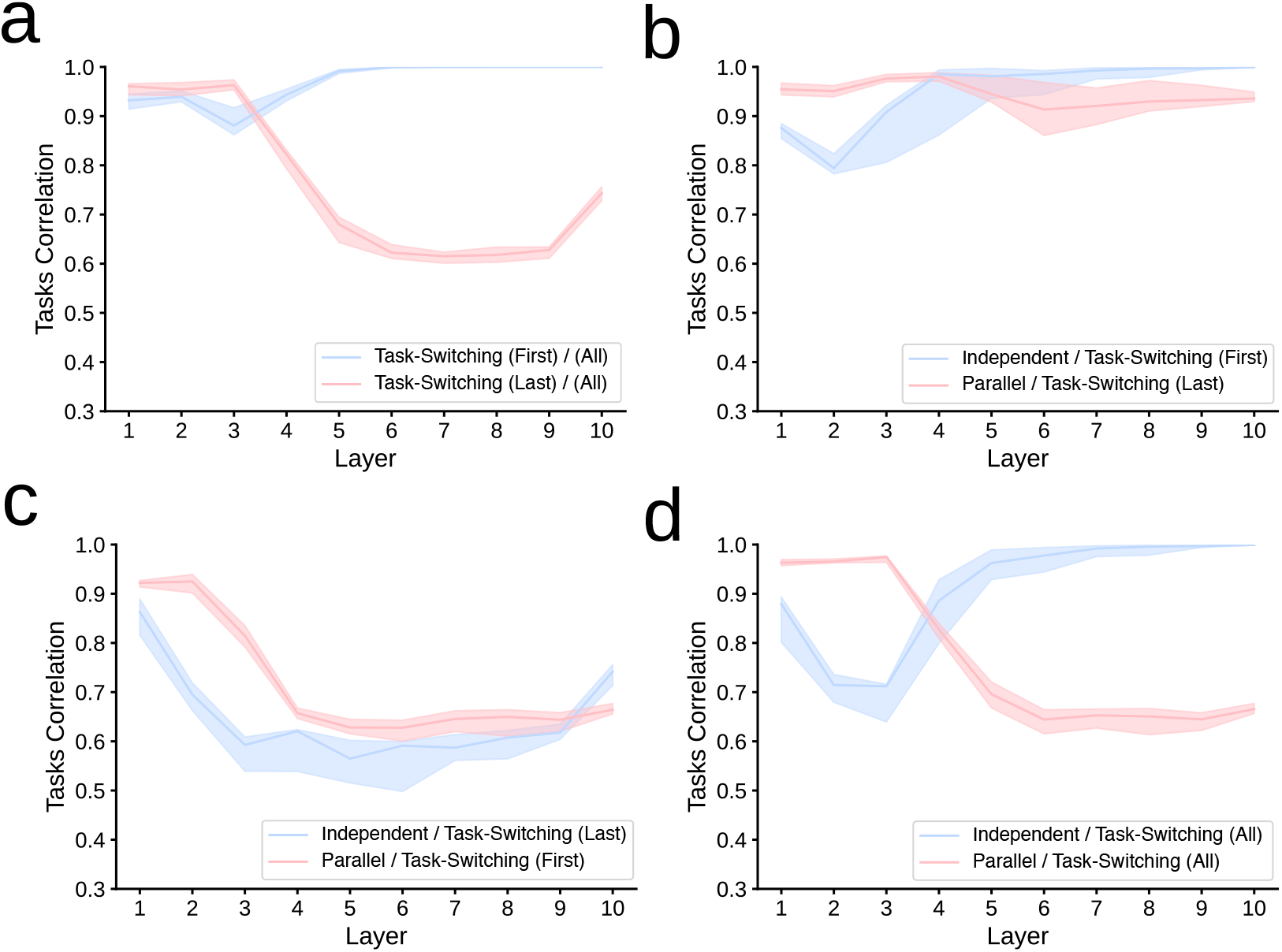
Task correlation between RDM’s main diagonal blocks. Pearson correlation coefficients between *independent, parallel* and *task-switching* networks across 10 different runs (Median and interquantile range, IQR, i.e., difference between the 75th and 25th percentiles of the data). In task-switching networks, the location of context inputs appears in parentheses: only in the first (First), only in the last (Last), or in every hidden layer (All).

Since the transition in Fig 3a, resembles that of Fig 2b, we compared *independent* and *parallel* networks with the *First* and *Last* layer networks, to quantify to what extent *First* and *Last* layers networks are representative of *independent* and *parallel* networks, respectively. We observe that task correlations between these models are very high (Fig 3b) in clear contrast to the opposite pairing (Fig 3c). These results suggest that the position of contexts determine the regime in which *task-switching* networks operate while performing multiple tasks. Contexts added only to the the first layer, drive representations similar to *independent* networks, whereas adding the contexts only in the last layer promotes *parallel* processing. Interestingly, *task-switching* networks with context in all layers (Fig 3d) promote *parallel* processing in early stages, to only transition to specific task processing thereafter, even though there are no location constraints for context inputs in the *All* layers model. Such *optimal* transition emerges from learning as the network converges to the best solution through training.

Results from a multidimensional scaling (MDS) analysis applied to RDMs (Jazayeri & Ostojic, 2021; Bernardi et al., 2020) (see details in Section Multidimensional scaling) further support this interpretation: adding context only in the first layer separates the representations of digits from both tasks, whereas postponing the context presence until the last layer, contrarily, prevents the separation of *parity* and *value* tasks (S1 Fig S6).

### Context impact is maximal at intermediate layers

Context inputs in *task-switching* networks can be seen as top-down inputs from PFC circuits mediating rule encoding in context-dependent decision making (Miller & Cohen, 2001; Menon & D’Esposito, 2022). The strength of the context may depend on the stage at which it is integrated into the sensory-response processing (Gilbert & Sigman, 2007). In many tasks, a progressive increase of attentional modulations in the visual cortex has been reported (Motter, 1993; McAdams & Maunsell, 1999).

In this section, we assess the strength and correlation of context inputs in the *task-switching* network (Fig 4). Previously, Flesch, Juechems, et al. (2022), observed anticorrelation between the task biases induced by context inputs in shallow networks with a single hidden layer. Here, we extend the analysis to check the impact of context inputs when the processing hierarchy is considered.

**Figure 4:**
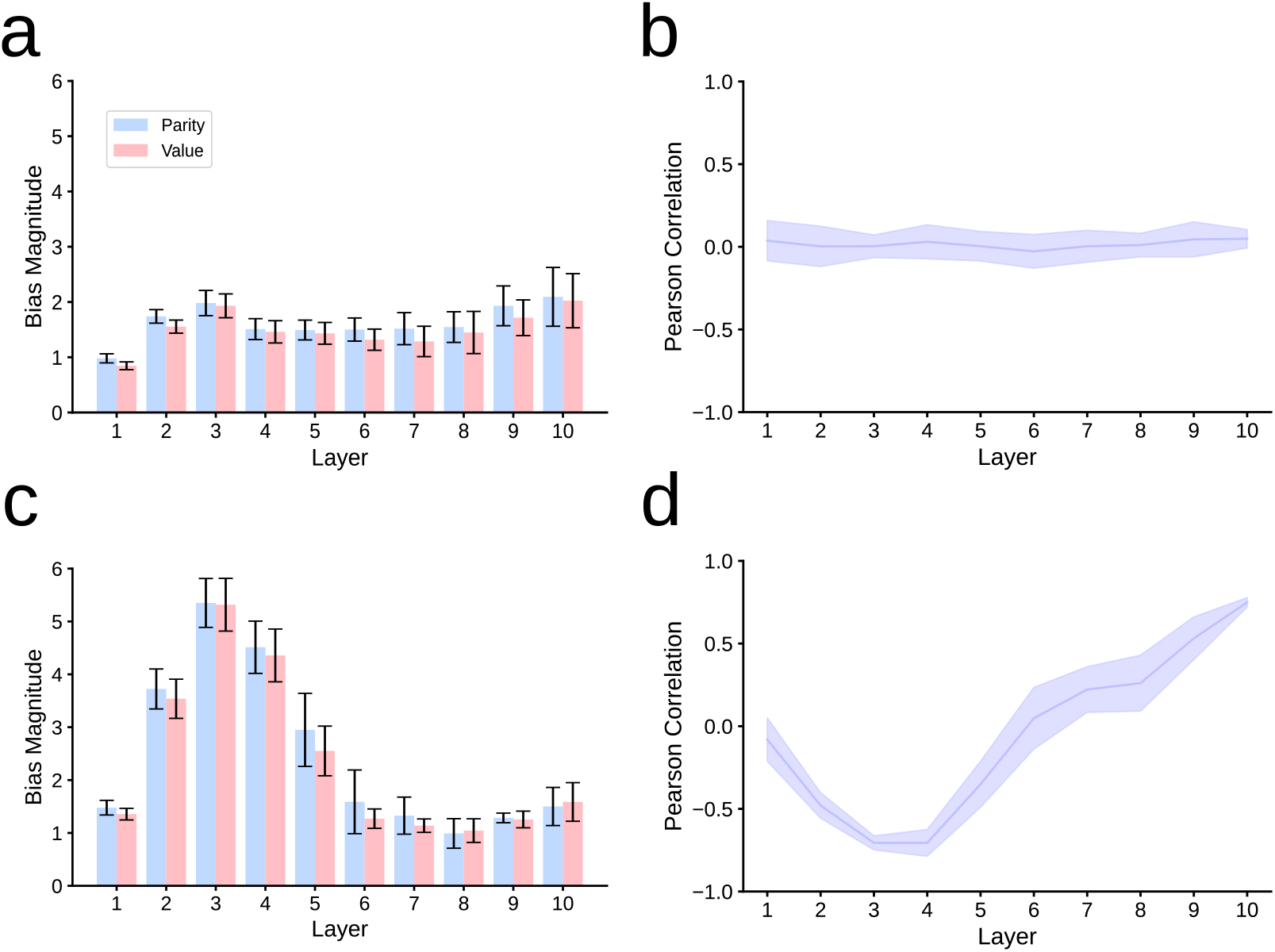
Bias connection strength and its correlation in *independent* and *task-switching* networks. Bias magnitude (a,c) and Pearson correlation (b,d) between input biases in *independent* networks (a,b), compared to *task-switching* networks (c,d) for the {parity, value} tasks (Mean and SD across 10 runs). Note that input biases are associated with context inputs only in *task-switching* networks. The magnitude of the bias is calculated using the *L*2-norm (see Section Context bias strength).

As control, we performed the analysis first in *independent* networks, for which bias parameters are task-agnostic. Indeed, for *independent* networks, the impact of the biases was not particularly sensitive to location neither in strength (Fig 4a) nor in correlation (Fig 4b).

In contrast, when analyzing the context inputs in *task-switching* networks trained for the same tasks, we observed that the connection strength increased importantly in the intermediate layers (Fig 4c), akin to advanced sensory processing in cortex. This increase in magnitude suggests an important role of context inputs at routing the sensory information toward the final output selection according to the task in play. This was further confirmed by the strong anticorrelation between the connection strengths of context inputs belonging to distinct tasks (Fig 4d). Analogous results were obtained for letter processing (S1 Fig S7. The distribution of connection strengths was very similar as well for digit processing with five tasks (S1 Fig S8).

We extended this analysis for *independent* and *task-switching* networks with different numbers of context and layers in S1 Fig S9, S1 Fig S10 and S1 Fig S11. Each row in these figures represents a different context configuration and each column denotes the index of the layer (we fixed the maximum number of layers to 10). Colored cells according to the heatmap refer to layers with context inputs at that position (S1 Fig S9, S1 Fig S10 and S1 Fig S11), whereas gray cells refer to layers with no context inputs (S1 Fig S9 and S1 Fig S10), and missing cells in S1 Fig S11 denote the absence of the layer, so networks were shallower. We tried four different settings: adding context inputs starting from the first layer (S1 Fig S9a), adding context inputs starting from the last layer (S1 Fig S9b), adding context inputs only at a particular layer (S1 Fig S10), and adding layers with context inputs progressively (S1 Fig S11). For all these conditions, we calculated the mean magnitude for each task (left and center panels) and the correlation between context-mediated task biases (right panels). Results from these analyses helped confirming that context inputs are stronger and more anticorrelated at mid-processing stages. Introducing context inputs only in a single layer results in even stronger magnitude and anticorrelation, regardless of the context location. Adding more layers did not affect the location where the magnitude of contexts and anticorrelation was the highest, which remained at intermediate layers.

### *Task-switching* networks encode task similarity

Experimental evidence suggests that neural coding in the brain may emerge by composition of simpler rules (Vaidya et al., 2021). In the PFC, the neural code used to store rule information over time is highly compositional (Reverberi et al., 2012). Neural network models extract task compositionality in paradigms designed by composition (Ito et al., 2022; Verbeke & Verguts, 2022). Since all parameters but those of contexts are shared in *task-switching networks*, context inputs are responsible for triggering the task processing.

In this section, we extend the correlation analysis between context connections across tasks (Fig 5a). We trained a ten-layer *task-switching* network in five different digit tasks. Some layers show only positive correlations (layer 1 and layers 7-10). Larger variability of pairwise task correlations is present in intermediate layers (layers 4-6), which include negative and positive correlations. We hypothesized that this variability might be associated with task similarity and influenced by the task congruency. To test this, we compared the pairwise context correlation matrix (that we will refer to as the similarity matrix), with the pairwise mean task congruency matrix, extracted from Table 2 after linearly scaling values from [0, 1] to [-1,1] (Fig 5b). The correlation between contexts in layer 4 was very tightly aligned with the scaled mean congruency between tasks.

**Table 1:** Task names and their output neuron encoding (labels).

| Task Name | Number |  |  |  |  |  |  |  |  |  |
| --- | --- | --- | --- | --- | --- | --- | --- | --- | --- | --- |
|  | 0 | 1 | 2 | 3 | 4 | 5 | 6 | 7 | 8 | 9 |
| Parity | 1 | 0 | 1 | 0 | 1 | 0 | 1 | 0 | 1 | 0 |
| Value | 1 | 1 | 1 | 1 | 1 | 0 | 0 | 0 | 0 | 0 |
| Prime | 0 | 0 | 1 | 1 | 0 | 1 | 0 | 1 | 0 | 0 |
| Fibonacci | 1 | 1 | 1 | 1 | 0 | 1 | 0 | 0 | 1 | 0 |
| Multiple of 3 | 0 | 0 | 0 | 1 | 0 | 0 | 1 | 0 | 0 | 1 |

**Table 2:** Pairwise mean task congruency across stimulus conditions.

|  | Parity | Value | Prime | Fibonacci | Multiple of 3 |
| --- | --- | --- | --- | --- | --- |
| Parity | 1 | 0.6 | 0.3 | 0.5 | 0.4 |
| Value | 0.6 | 1 | 0.5 | 0.7 | 0.4 |
| Prime | 0.3 | 0.5 | 1 | 0.6 | 0.5 |
| Fibonacci | 0.5 | 0.7 | 0.6 | 1 | 0.3 |
| Multiple of 3 | 0.4 | 0.4 | 0.5 | 0.3 | 1 |

**Figure 5:**
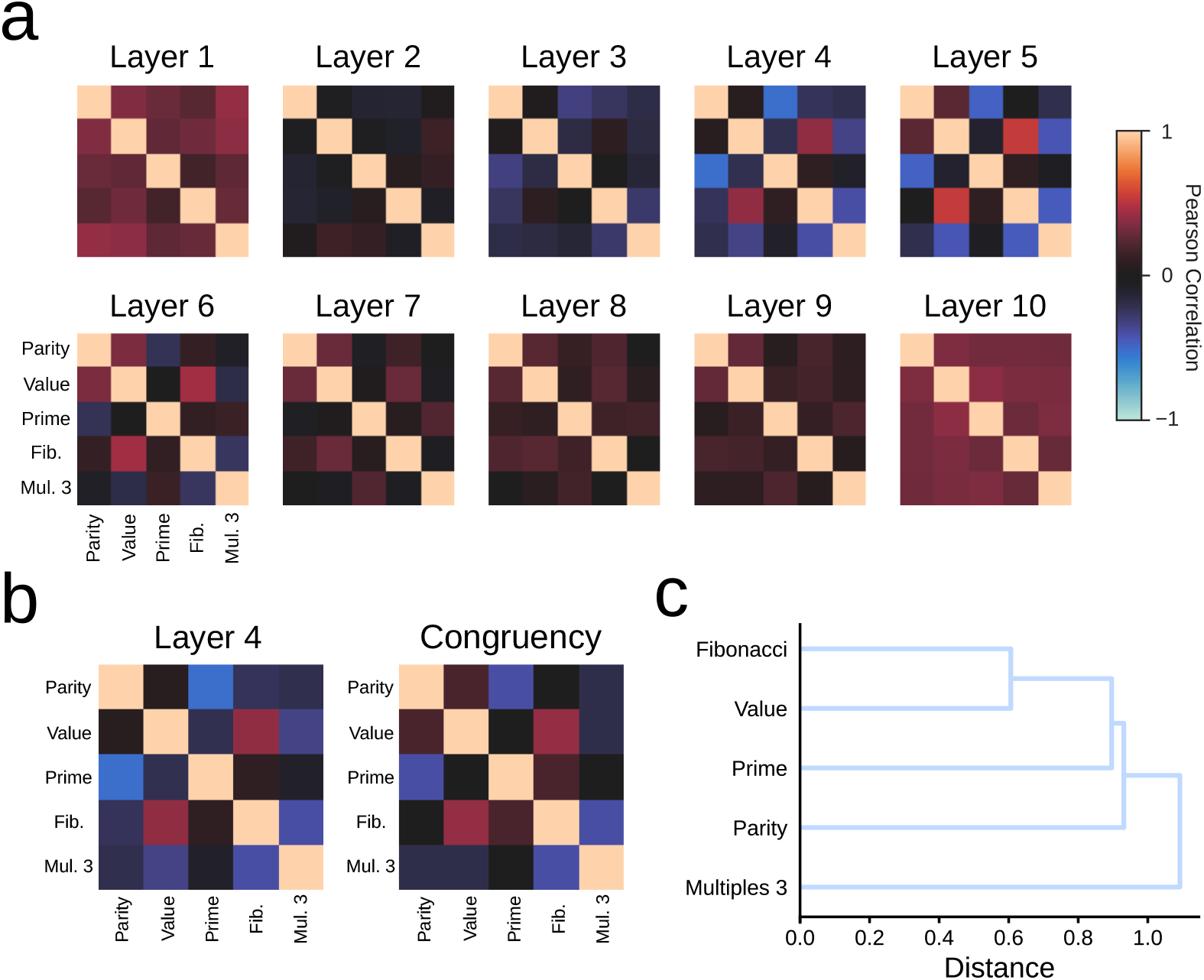
Average pairwise correlation between context connections for 5 distinct digit tasks, {*parity, value, prime, fibonacci* and *multiples of 3*}. (a) Task similarity matrices computed with the Pearson correlation across layers (legend on the right). (b) The similarity between contexts in layer 4 (left panel) vs. the pairwise mean task congruency linearly scaled to the [-1,1] range (right panel). (c) Dendrogram of task similarity in layer 4.

We quantified this further by computing the pairwise correlation between the scaled mean task congruency and the similarity matrix between context connections across layers (S1 Fig S12). The highest correlation occurs in layers 3–5 (S1 Fig S12), coinciding with the layers where the strengths of context connections were the largest (S1 Fig S8). Thus, in the sequence of internal processing, *task-switching* networks learn through training to first extract the mean task congruency in intermediate layers (S1 Fig S12). From there, *task-switching* networks unwrap the task similarity making shared representations grow toward the final goal (see Fig 2c, S1 Fig S1 and S1 Fig S2).

We further assessed the similarity between tasks from the similarity matrix in layer 4, using three methods: firstly, we computed a dendrogram (Sun et al., 2021), in which distance was represented by dissimilarity (1 − *r*, where *r* represents the correlation coefficient, Fig 5c); secondly, we applied multidimensional scaling to create a 2D low-dimensional projection of the similarity matrix (S1 Fig S13); finally, we trained a linear decoder to study how layers extracted features along the processing in the network (Alain & Bengio, 2016; Kriegeskorte & Douglas, 2019) (S1 Fig S14). The latter analysis reveals that *independent* networks primarily encode the task output regardless the layer (S1 Fig S14, mid panels), while *parallel* networks primarily encode congruency (S1 Fig S14, right panels). In contrast, *task-switching* networks are versatile. The model starts classifying the stimulus by type (S1 Fig S14, left panels), then groups stimuli according to congruency (S1 Fig S14, right panels), and finally maps them to the right task output (S1 Fig S14, mid panels). These results also support the hypothesis that *task-switching* networks learn to extract congruency information between tasks as an intermediate process toward the task goal.

### *Task-switching* networks are robust against *physiological* context modulations

In *Task-switching* networks, complimentary binary contexts 0/1 specify the task to perform. After training, we can modulate the strength of context inputs to analyze its impact in neural representations and network performance, akin to the strength of task signals in the PFC and their impact on task-relevant information processing (Egner & Hirsch, 2005).

First, we interpolated the values of context inputs between [0, 1] for the two tasks keeping their sum to 1 (see methods Section Context bias interpolation). We can make an analogy for context values in the ]0, 1[ range, so that values take into account the conflict between competing tasks (Fischer & Plessow, 2015), or the uncertainty in the context-encoding neural circuit, reinforced for instance by probabilistic paradigms. Results show that the transition between tasks (from *value* to *parity*) was smooth (Fig 6). There is a progressive change from one task to the other without an abrupt transition. The neural representations at 0.5 showed an intermediate state between tasks, where the accuracy (0.8) was at chance level, once the congruency baseline (0.6) was considered.

**Figure 6:**
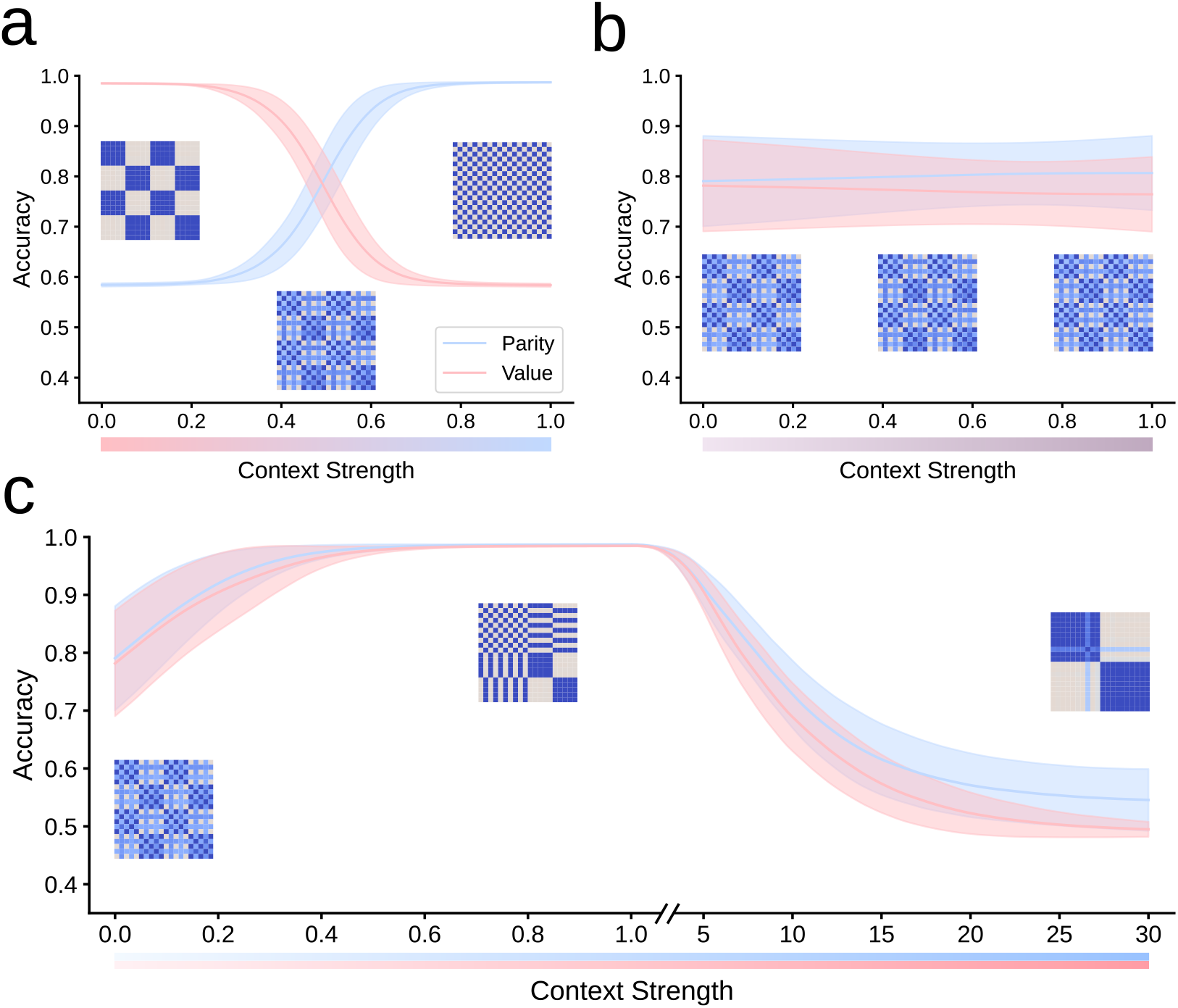
Impact of context strength on accuracy. (a) The total context value is fixed to 1. (b) Covariation of context values. (c) Independent contexts inputs of increasing value, while the other context remains at 0. Mean accuracy and SD across 10 different runs is shown, as well as RDM insets in different scenarios.

We then incremented the value of both contexts together from 0 to 1 (Fig 6b). This is a way to parametrically analyze performance according to context strength when both tasks are equally likely. The accuracy stayed close to chance level (0.8), regardless of the context strength. The internal representations, displayed in the RDM panels, show that the network operates midway between the two tasks, generalizing the observation in Fig 6a, when both tasks had the context strength equal to 0.5.

Lastly, we analyzed the case when one context strength was varied keeping the other at zero, and tested for values lower and greater than one (Fig 6c). Values lower than one can be related to weak context modulations, whereas values greater than one are useful to infer behavior against aberrant context inputs, such as those that could be originated from abnormal activity in the prefrontal cortex. Results indicate that networks are robust against fluctuations in context strength in a wide range. Concretely, we observed that even though the strength of independent contexts was reduced to 0.5, or enhanced to 5, the network was still reaching an accuracy close to optimal. Beyond those points, however, performance was significantly impaired (Fig 6c). When the context dramatically increased, the network was unable to process the stimulus (digit) upon which select the task output, which led to a decrease in accuracy. This occurred because the information about the task to perform masked the digit identity, and both are needed for proper action selection.

These results align with a neuroscience perspective of context-dependent decision making: task performance depends on how the context maps the sensory information to the appropriate response. In general, this mapping depends on (a) context uncertainty: biased competition between rule encoding cells in prefrontal cortex unbalance the probability to respond according to either task; and on (b) context strength, akin to the strength of top-down modulations, according to which we can observe three distinct scenarios: when the context strength is abnormally weak, the task to perform is uncertain, and performance collapses (Papyan et al., 2020) to a chance level based on the mean task congruency (Fig 6c, left side); when the context strength is within a physiological range, performance is close to optimal for all tasks (Fig 6c, center); and when context strength is abnormally high, performance is dramatically impaired approaching a 1*/N* chance level, where *N* represents the number of alternative responses (Fig 6c, right side). In this latter scenario, there is complete certainty about the task to perform, yet the context input is so strong that masks the sensory information, resulting an equal probability to select either output response.

### Neural selectivity emerges to resolve task execution

Distinct neural assemblies in the brain coordinate to process information and shape neurons’ selectivity (Hebb, 1949). There is evidence in favor of subnetworks acting as units of processing, which is known as the neural population doctrine (Yuste, 2015; Saxena & Cunningham, 2019; Ebitz & Hayden, 2021).

We hypothesized that similar mechanisms operate in *task-switching* networks, so that neurons may be grouped by contextual information into different subnetworks. Clustering techniques upon neural data can be utilized to functionally characterize group of neurons that act similarly (Ardid et al., 2015). To test our hypothesis and investigate the emergence of subnetworks in *task-switching* networks, we clustered neural activations. The goal of the analysis was to characterize different neuron assemblies according to the stimulus input and task context (see details in Section Clustering of neural activations).

Fig 7 shows the mean activation of neurons in the *parity* and *value* tasks for the two clusters when presented each one of the digit inputs. The *removed* panel was obtained when no context information was introduced to the *task-switching* network. Results from this analysis identifies two clusters, or neural assemblies. For the *parity* task, we found a cluster encoding even digits (in blue) and another cluster enconding odd digits (in red). And similarly for the *value* task, encoding values smaller than or equal to four (in blue), or larger values (in red). The congruent digits (same output for the two tasks) showed the most salient activations. In fact, when context inputs were removed, neural assemblies collapsed, encoding only the congruent digits (Papyan et al., 2020), and entailing that context inputs are responsible for the increase in activation for incongruent stimuli. In addition to the congruent/incongruent asymmetry, the patterns of activation also showed an asymmetry between output classes: in the *parity* task, the activations to even digits were dominant against odd digits, and similarly in the *value* task for the activations to values smaller than or equal to four against larger values. The predominant output classes varied randomly for different runs.

**Figure 7:**
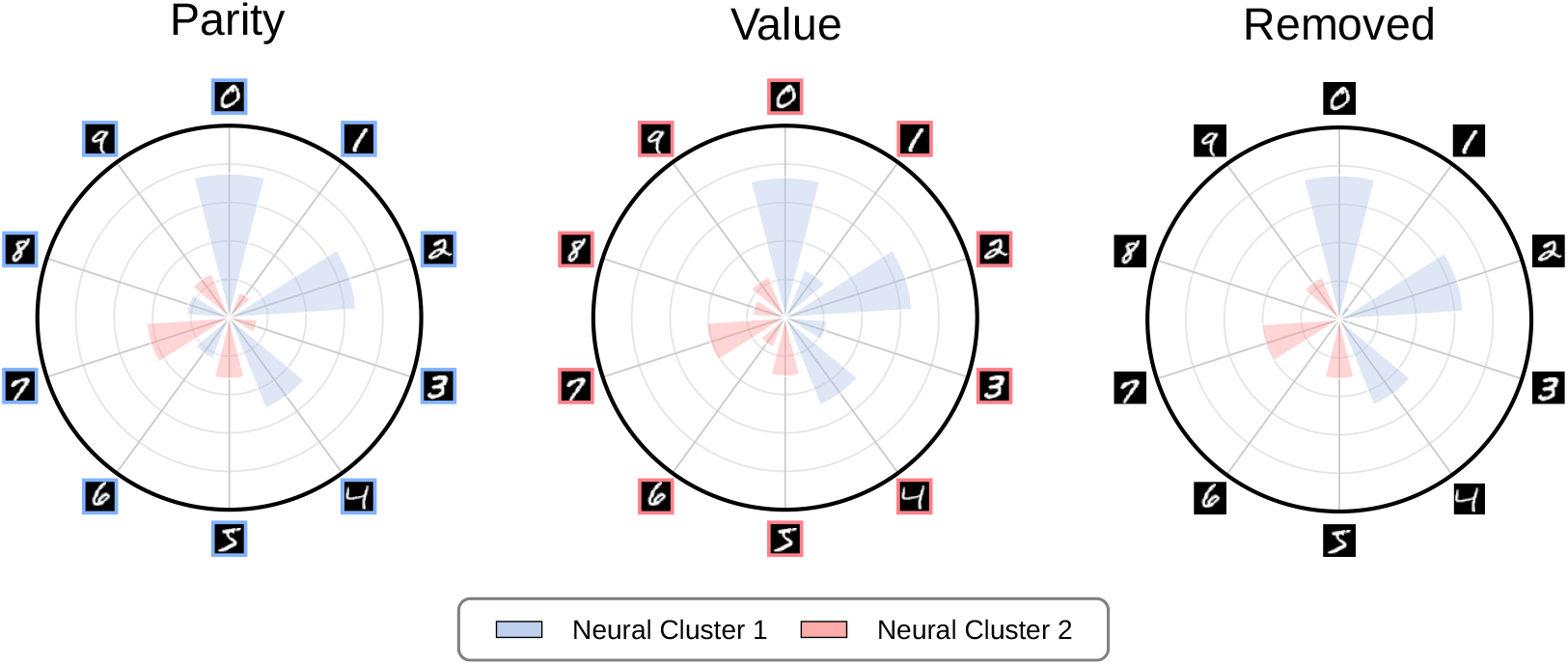
Neural clustering for the {*parity, value, removed*} *task-switching* schedule. *Removed* denoted the condition in which no contextual information was introduced.

We also repeated the analysis using five digit tasks (S1 Fig S15). In this analysis, we observe that the two different neural assemblies encode different digits depending on task context, showing that ensembles in *task-switching* networks can be tuned to the interaction of stimulus and task identities.

## Discussion

In this study, we conducted a series of analyses with multitask networks, namely *independent, parallel* and *task-switching* networks. We studied these networks using representational similarity analysis (RSA) (Kriegeskorte et al., 2008; Kriegeskorte & Kievit, 2013), a methodology developed in neuroscience to relate task and stimuli with internal representations (Fig 2). We defined shared representations as a function of the off-diagonal of the representational dissimilarity matrices (RDM). The amount of shared representations of a network depends on the architecture and the tasks to be learned. For instance, *independent* networks have low shared representation by design, since there is no parameter sharing between task modules. In the case of *task-switching* and *parallel* networks, shared representations are related to the generalized congruency between tasks. The *generalized congruency* is a powerful concept that we introduce in this manuscript as an extension of the traditional *stimulus congruency* and *categorization* definitions. Upon the generalized congruency, we were able to predict the final representation motifs of *task-switching* and *parallel* networks. This theoretical approach help distinguishing the final representational convergence of the two networks: single task processing (main diagonal) and pairwise task processing (off-diagonal) in the RDM of *task-switching* networks, regardless of the number of tasks (S1 Fig S19). In contrast, the final motif in *parallel* networks represent multitask processing and vary with the number of tasks. Increasing the number of tasks, degrade the RDM patterns of activation in *parallel* networks, affecting their interpretability by visual inspection, even though our theoretical approach explains the final motif for *parallel* networks as a linear combination of tasks *union* and *intersection* (S1 Fig S20). This discrepancy between the two architectures is reflected on diminished shared representations for *parallel* in comparison to *task-switching* networks.

Context inputs facilitate feature extraction in alignment with the task they encode, but their impact depends on their position in network’s processing. When context is only added in the last hidden layer, *task-switching* networks mostly operate as *parallel* networks, since there is no context information beforehand to route activity toward the task. On the other hand, training *task-switching* networks with the context input only in the first hidden layer precludes the extraction of common task features, as it happens in *independent* networks (Fig 3). Our results suggest that *independent* and *parallel* networks can be seen as particular *task-switching* networks, if contexts are introduced only in early processing stages (*independent* networks), or if context inputs are removed or delayed (*parallel* networks). This picture also emerges when contexts are continuously varied in the [0, 1] interval: the pattern of *parallel* shared representations appear for diminished context inputs, whereas the pattern of *independent* representations emerge as context inputs grow (Fig 6c).

We estimated the importance of context biases in the network processing by analyzing the magnitude of their connections at different processing stages. Biases with higher magnitude affect neurons in greater measure. Hence, a way to activate different downstream subnetworks is through biases having dissimilar impact upon distinct neurons. In artificial neural networks, mechanisms for activating different subnetworks to perform different tasks have been previously studied (Rosenbaum et al., 2017; Wortsman et al., 2020). Our results show that intermediate layers have greater importance at modulating downstream activation (Fig 4b, S1 Fig S9 top panel and S1 Fig S11). Moreover, the highest anticorrelation between context connections also occurs in the same layers (Fig 4b, S1 Fig S9 top panel and S1 Fig S11), supporting the mechanism. We also investigated the correlation between context connections after learning a larger amount of tasks. This analysis informed that the layers with greatest magnitude learn task similarity by encoding the pairwise mean task congruency (Fig 5). The interpretation was also supported after training a linear decoder to study how layers extracted features along the processing in the network (Alain & Bengio, 2016; Kriegeskorte & Douglas, 2019) (S1 Fig S14).

Lastly, we analyzed how learning shaped neurons’ selectivity in *task-switching* networks. We observed that neurons in the last layer clustered together according to the task and the output response (Fig 7). These results suggest that the integration of the stimulus and context information drive the activation of different subnetworks, promoting the selectivity of distinct cell assemblies. Further work could investigate if single cells in the assemblies present non-linear mixed selectivity (Rigotti et al., 2013), argued to promote reliable coding strategies in the presence of noise (W. J. Johnston et al., 2020), and to increase the dimensionality of neural representation, suggested to facilitate linear decoding by downstream neurons (Fusi et al., 2016).

Our work shows that context biases are able to guide goal-directed behavior in simple feedforward neural networks, as they do in brain circuits of living agents. For the sake of explainability, applying our theoretical approach and extending our study of neural representations to more sophisticated biological and artificial systems may help shedding light on the internal processing of complex architectures (Samek & Müller, 2019). In deep neural networks, the usage of contextual information has been explored in terms of fixed embeddings: task embeddings for multitasking (Li et al., 2016; Liu et al., 2019; Sun et al., 2021), class embeddings for conditional data generation (Brock et al., 2018), and embeddings for text-to-image generation (Radford et al., 2021). Further research could focus on the analysis of internal neural representations, to understanding the impact of fixed embeddings in comparison to context biases. Remarkably, context inputs have allowed networks to learn tasks sequentially (Masse et al., 2018; Serra et al., 2018; Grewal et al., 2021; Flesch, Nagy, et al., 2022), without forgetting previously learned tasks, hence avoiding catastrophic forgetting (French, 1999). The characterization of neural representations in complex architectures, however, is still pending, and our approach could set the basis for such studies.

## Acknowledgements

This work was supported by the Generalitat Valenciana Gen-T Program (Ref. CIDEGENT/2019/043), Grant Ref. PID2020-120037GA-I00 funded by MCIN/AEI/10.13039/501100011, and European Union NextGenerationEU/PRTR Programa de Planes Complementarios I+D+i (Ref. ASFAE/2022/014).

## Methods

## Materials and methods

### Neural network architectures

We conducted a set of analyses with three variants of a feedforward network (Rumelhart et al., 1986; Goodfellow et al., 2016) in the context of multitask learning (Fig 1).

- *Parallel Network*: All tasks 𝕋 = {*t*_1_, *t*_2_…*t*_*T*_ }, assuming *T* tasks, are processed identically in hidden layers. Only the output layer accounts for task-independent responses. The output of each hidden layer in a *parallel* network is parameterized by **y** = *f* (**W**^*T*^ **x** + **b**), where **x** denotes the inputs to the layer, *f* denotes the activation function, and **W** and **b** denote respectively shared synaptic weights and biases for all tasks. The only independent parameters appear in the output layer, which is parameterized by 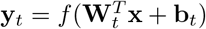, where *t* denotes a specific task in the 𝕋 task set.
- *Independent Network*: Each task is learned in isolation through unconnected subnetworks. Multitasking is performed by combining the independent outputs. The output of each layer in a *independent* network is parameterized by 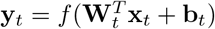 for each task *t* in 𝕋. Neither inputs, nor weights and bias parameters are shared across tasks since *independent* subnetworks are mutually disconnected.
- *Task-switching Network*: Each task *t* in T is processed individually, even though parameters other than the context biases are fully shared. The output of each layer in a *task-switching* network is parameterized by **y**_*t*_ = *f* (**W**^*T*^ **x**_*t*_ + **b**_*t*_). The same weights apply to all tasks, whereas context biases **b**_*t*_ are task specific.

*Task-switching* networks require at least one layer with the context biases. We consider cases in which the context information is not presented to a subset of layers. For those, the output of layers before the introduction of contexts is not task specific **y** = *f* (**W**^*T*^ **x**), whereas the output for layers following the introduction of contexts inherit task information only through their inputs **x**_*t*_: **y**_*t*_ = *f* (**W**^*T*^ **x**_*t*_). Mathematically, the former case is akin to the expression for hidden layers in *parallel* networks with the exception of missing the bias parameter, whereas the latter case shares with *independent* networks only the fact that the input is task specific. Nevertheless, we will show in results that these similarities suffice to replicate in the *task-switching* networks the internal representations of *parallel* and *independent* networks.

### Context bias strength

The strength of the context biases in each hidden layer **b** ∈ ℝ^*H*^, where *H* represents the number of neurons in the layer, is normalized by ||**b**|| according to the *L*2-norm:

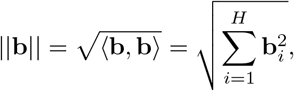

where **b**_*i*_ represents the context bias to neuron *i*.

### Context bias interpolation

Given two context biases **b**_1_ and **b**_2_, we can find a bias 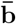 which is a linear combination **b**_1_ and **b**_2_:

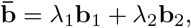

By adding the constraint *λ*_1_ + *λ*_2_ = 1, we can interpolate from **b**_1_ to **b**_2_ by sweeping *λ*_1_:

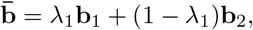

or from **b**_2_ to **b**_1_ by sweeping *λ*_2_:

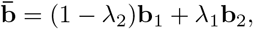

We used this approach to interpolate the values of context inputs between [0, 1] for the two tasks in Fig 6, keeping their sum to 1.

### Two-alternative choice tasks

In this study, we considered a *two-alternative choice tasks* using digits with the MNIST dataset (LeCun et al., 1998), and lowercase letters using the EMNIST dataset (G. Cohen et al., 2017). We mainly used the *parity* task (even vs. odd digits) and the *value* task (digits larger than four vs. not), which have been previously used in human experiments analyzing task switching (Monsell, 2003), and in neural networks analyzing multitasking (Bernardi et al., 2020).

To test the generality of our results, we also analyzed *two-alternative choice tasks* in the domain of letters, the *vowel* task (vowel vs. consonant) and the *position* task (first vs. second half of the alphabet).

In addition, to analyze task scalability, we designed up to five distinct digit tasks using the MNIST dataset. Assuming that the performance was independent among tasks, the joint accuracy for *T* tasks was inferred from the product of the probabilities to perform well in each individual task:

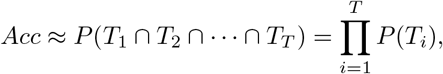

We also analyzed the impact of *congruency* in learning representations in the *task-switching* network. Table 1 enumerates the tasks and the expected output neuron of the network for digit tasks, i.e., the labels (0, for the first output neuron, and 1 for the second output neuron).

A stimulus is defined *congruent* for two or more tasks if the same output is selected for those tasks. For example, the number 0 is congruent for *parity* and *value*. The opposite, *incongruency* of a stimulus occurs when the output selection to a stimulus is different for the distinct tasks. Following the example, the number 6 is incongruent for *parity* and *value*. The congruency ratio between tasks is represented in Table 2.

In the two-alternative choice tasks with letters, we define vowels and the first half of the alphabet to take label 1, whereas consonants and the second half of the alphabet take label 0. A congruent letter for *vowel* and *position* would correspond, for instance, to the letter *a*. An incongruent letter would be, for example, letter *b*.

Strictly speaking, congruency applies to *task-switching* networks but not to *independent* and *parallel* networks since these networks do not share the same output neurons across tasks. However, for the sake of network processing interpretation, we generalize the concept of congruency in *independent* and *parallel* networks by keeping the output labeling of *task-swiching* networks across tasks (see Table 1).

Note also that the congruency does not directly imply task dependence: the probability to select an output neuron varies for each individual task even for a congruent stimulus to these tasks. Thus, the probability to select the correct output neuron for the 0 digit in the *parity* and *value* tasks is different, which is caused by the modulation of distinct context biases.

### Training

The total number of images in MNIST is 60,000, evenly distributed among digits. We split MNIST into a training set of 50,000 images and a test set of 10,000. For EMNIST, we used only a subset to guarantee that every lowercase letter had the same frequency. The total number of images in EMNIST was 49,296, with 1896 images for each letter. We split EMNIST into a training set of 41,080 images and a test set of 8216. For both datasets, we fixed the batch size to 100.

The models were implemented in PyTorch (Paszke et al., 2019). For each architecture, we trained two different models with 5 (Section *Task-switching* networks operate in a midway regime between *independent* and *parallel* networks) and 10 layers (Sections Context location determines the transition between *independent* and *parallel* processing, Context impact is maximal at intermediate layers, *Task-switching* networks encode task similarity, *Task-switching* networks are robust against *physiological* context modulations, Neural selectivity emerges to resolve task execution). For the *task-switching* network, we followed a interleaved shuffled schedule, where tasks were presented in random order during training (Flesch et al., 2018).

Networks were trained for 50 epochs. The number of hidden neurons per layer was fixed to 100. We used ReLU as the activation function (Glorot et al., 2011) and initialized weights and biases following a uniform distribution in the range 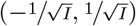, where *I* is the number of inputs to the layer. We used the Adam optimizer (Kingma & Ba, 2014) with the learning rate set to 0.001, *β*_1_ = 0.9 and *β*_2_ = 0.999, and selected cross-entropy as the loss function. We collected statistics over 10 different initializations (*runs*) for hypothesis testing, controlling the random number generator for reproducibility. We use the Wilcoxon Signed-Rank Test to test statistical significance between models across runs (Demšar, 2006).

## Representational dissimilarity matrix

Representational dissimilarity matrices (RDMs) capture neurons’ coactivation (Kriegeskorte et al., 2008). We analyze RDMs for each hidden layer to quantify the dissimilarity of internal representations across task conditions and input stimuli, such as digits or letters.

### Construction of RDMs

For each hidden layer, we construct an activation matrix of dimension *H*×*T*×*S*, where *H* runs over each neuron, *T* is the number of tasks, and *S* represents the number of stimulus types (digits or letters) in the test set. For each neuron, task and digit stimulus in 0–9 (or letter in *a*–*z*), we calculate the average neuron activation across the individual stimuli items (e.g., distinct handwritten 0’s). We then compute the correlation matrix **R** of dimension *TS* × *TS*, where *R*_*ij*_ denotes the pairwise correlation between average activation vectors 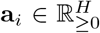 and 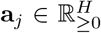. With these constraints, *R*_*ij*_ ∈ [0, 1] for cosyne similarity, and *R*_*ij*_ ∈ [−1, 1] for Pearson correlation.

The RDM is then calculated as:

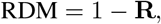

which is organized concatenating first by task, then by input type. For Pearson correlation, the RDM is bounded in the range [0, 2], 0 representing complete similarity (correlation) and 2 complete dissimilarity (anticorrelation). For cosyne similarity, the RDM is bounded in [0, 1], since all activations are ∈ℝ _≥0_. This is because the ReLU activation function is used in the models in all hidden layers.

For *independent* and *task-switching* networks, the RDM can be computed directly. For *independent* networks, we have specific activity patterns from each network performing each task, and for *task-switching* networks, we have specific activity patterns originated from the same network under each task context. This is, however, not the case for *parallel* networks, since internal representations for both tasks are identical. For the sake of easing the comparison among models, the RDM of *parallel* networks is composed following tile concatenation from the same activation patterns.

### Sensitivity to data variability

When constructing RDMs, we average the activations across inputs of the same type, such as different instances of the 0 digit or the a letter. Since the accuracy of the networks surpasses 90% accuracy for the tasks, the noise introduced by misclassification is negligible. S1 Fig S17 shows the comparison between RDMs with and without averaging.

To test the robustness of RDMs to data variability, we have added random gaussian noise parameterized by its standard deviation *σ* to test the degradation in the RDMs. We calculate the signal-to-noise ratio (SNR):

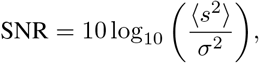

where ⟨*s*^2^⟩, denotes the average signal power (computed from the pixels’ intensity in the image), and *σ*^2^ is the noise variability. Results of this analysis (S1 Fig S18) show that even when the noise variability is of greater strength than the signal power (for example for *σ* = 0.5), noisy RDMs highly maintain the structure of noiseless RDMs. Yet, if noise variability is further increased (*σ* = 1), RDMs become harder to interpret visually, even though they significantly correlate with noiseless RDMs.

### Shared representations

We estimated shared representations from RDMs. The RDMs are conformed by submatrices or blocks of size *T* × *T*, where *T* runs over tasks. This conforms two different regions, the main diagonal and the off-diagonal.

The total number of blocks of the RDM is *T* + 2*O*_*D*_, where *T* is the sum of the blocks in the main diagonal, which equals to the number of tasks, and *O*_*D*_ includes the blocks in either the upper or the lower off-diagonal.

We estimated shared representations (SR) from the Pearson correlation between activation vectors belonging to the off-diagonal blocks.

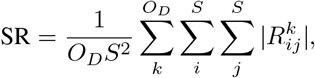

where *O*_*D*_ is given by 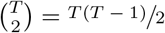, and 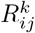 corresponds to the correlation at position *ij* of block *k*.

For completeness, task representations (TR) can be analogously computed repeating the approach over the blocks in the main diagonal:

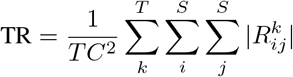

The complete representation (CR) of the RDM corresponds to:

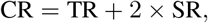

since the number of blocks is *T* + 2*O*_*D*_ and the RDM matrix is symmetric.

### Generalized congruency

The concept of congruency is applied to specific stimuli, so that a stimulus is defined *congruent* (or *incongruent*) for two or more tasks if the same output (or different) is selected for those tasks. In this view, congruency separates stimuli into congruent and incongruent subclasses. Consequently, the concept is frequently referred to as *stimulus congruency*.

We realized that generalizing the definition of congruency, setting tasks and stimuli at the same level, is important to better interpret network internal representations. In this view, the *generalized congruency* is defined in terms of the same vs. different output for the two distinct conditions. Thus, if the output for task 1, when presented the *i* stimulus, is the same as the output for task 2, when presented the *j* stimulus, the two conditions are congruent; otherwise they are incongruent.

To better see this, we can examine a couple of generalized congruency conditions: number 0 for the *parity* task and number 3 for the *value* task are congruent, since these two conditions map to the same output label; whereas number 0 for *parity* and number 6 for *value* are incongruent, since these other two conditions map to either output label (see Table 1).

In fact, the *generalized congruency* definition reduces to the *stimulus cogruency* if, between the two compared conditions, the same stimulus is considered. Remarkably, the generalized congruency also supersedes the concept of *categorization*, defined as the set of stimuli that, in the context of a given task, map to the same output, hence belonging to the same class. Thus, if we restrict the generalized congruency to a specific task, its definition reduces exactly to that of *categorization*.

In the next section (Final representation motifs), we show that final representations in RDMs can be predicted from the generalized congruency. It reduces to the categorization mapping of each task in the RDMs’ main diagonal, whereas the resulting mapping for each given stimulus in RDM’s off-diagonal sets its (stimulus) congruency. Beyond those two special cases, final representations also inform about distinct stimuli under distinct tasks pointing to the same response.

### Final representation motifs

Representation block motifs in the RDMs for the last hidden layer of the *task-switching* network can be predicted from the generalized congruency of tasks with binary outputs: {0, 1}.

To see this, we first introduce a function *f* that computes the pairwise mean task congruency, that is fraction of congruent stimuli for two tasks. Then, we construct, by analogy, a function *F* that computes the generalized congruency, which predicts the last hidden layer (i.e., final) representation motifs in the structure of RDMs for *task switching* networks. Finally, we describe how representation final motifs in *task-switching* networks relate to those of *independent* and *parallel* networks.

Let’s begin considering two tasks expressed as column vectors of dimension *S*: **t**_1_ ∈ {0, 1} ^*S*^ and **t**_2_ ∈ {0, 1} ^*S*^, where *S* runs over all stimulus conditions (e.g., 0–9, for digits). We can define a function *f* (**t**_1_, **t**_2_) that maps the two task vectors to a scalar representing the fraction of congruent stimuli 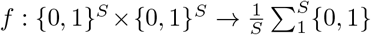 (see Table 2). Given the binary nature of the outputs, we can express *f* as:

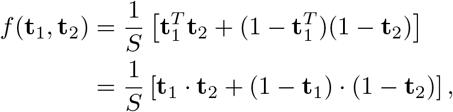

where *T* is the transpose operator and · denotes the dot (or inner) product. Mathematically, *f* = 1 when **t**_1_ = **t**_2_, meaning that all stimuli are congruent for a single task. If we consider, for instance, that **t**_1_ and **t**_2_ represent the *parity* and *value* tasks, respectively, then **t**_1_ = [1, 0, 1, 0, 1, 0, 1, 0, 1, 0]^*T*^ and **t**_2_ = [1, 1, 1, 1, 1, 0, 0, 0, 0, 0]^*T*^ (see Table 1), and the resulting fraction of congruent stimuli is *f* (**t**_1_, **t**_2_) = 0.6 (see Table 2).

Similarly, we can define a second function *F* : {0, 1}^*S*^ ×{0, 1}^*S*^ → {0, 1}^*S*×*S*^, that maps the two task vectors into the square *S* × *S* generalized congruency matrix as follows:

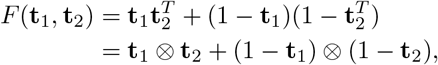

where ⊗ denotes the outer product. The entry *F*_*ij*_ will be equal to 1 only if the output for the two tasks is the same *t*_1_(*i*) = *t*_2_(*j*), i.e., if the output for task 1, when presented the *i* stimulus, is the same as the output for task 2, when presented the *j* stimulus; otherwise *F*_*ij*_ will be equal to 0 (see Section Generalized congruency). If tasks differ, the resulting matrix *F* (**t**_1_, **t**_2_) is not symmetric.

From *F*, we can compute the mean value of each final block motif for two tasks, **t**_1_ and **t**_2_, as follows:

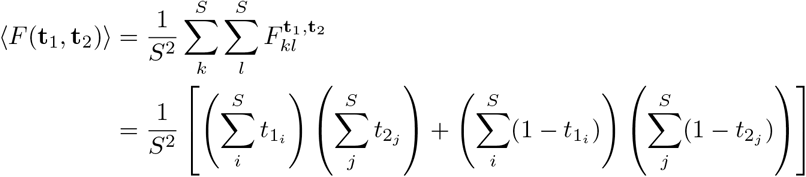

We can generalize the approach for an arbitrary set of tasks 𝕋 = {**t**_1_, **t**_2_…**t**_*T*_ } with *T* tasks:

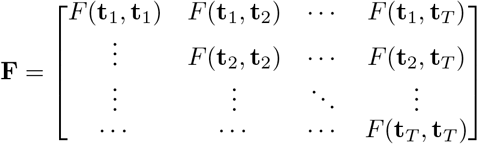

The generalized congruency matrix *F* can be compared to the final representation motifs of the RDM. Given that each term in *F* is bounded to {0, 1}, computing the empirical RDM in terms of cosyne similarity, compared to Pearson correlation, provides a more precise relationship with the theoretical approach.

S1 Fig S19 compares the empirical final block motifs of the RDM in a 10-layer *task-switching* network with its prediction. For the prediction, we have assumed that the correlation matrix **R** converges to **F** in the final representation (i.e., that of the last hidden layer). Thus, the *task-switching* network converges to a RDM of the form:

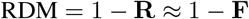

Furthermore, we can estimate the amount of shared representation in *task-switching* networks by averaging the mean value of block motifs located in the off-diagonal of the RDM matrix (see Section Shared representations):

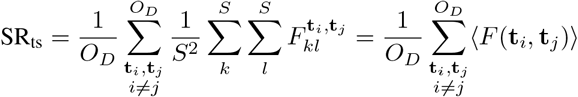

S1 Fig S19 also compares the predicted mean similarity representation with that in the empirical final block motifs of the RDM, showing a good estimation of shared and task representations.

The same approach can be applied to the *independent* and *parallel* networks. For the *independent* network, the final motifs of the main diagonal are identical to those in the *task-switching* network. However, the motifs in the off-diagonal differ (see Fig 2a) because the activations of neurons belonging to disconnected subnetworks in the *independent* network are uncorrelated.

Note that, by construction, there cannot be shared representations in the *independent* network. Fig 2c, however, displays residual shared representations, which come from the fact that activations covary just by chance for a fraction of pairwise neurons in the independent subnetworks. This residual level in Fig 2c can be interpreted as the chance level in the estimation of significant shared representations. Because of that, we computed RDMs and estimated shared representations based on Pearson correlation since this was the analysis that best controlled residual correlations in comparison to cosyne similarity and Spearman correlation. As mentioned above, we limited the use of cosyne similarity only to the theoretical approach developed in the current section.

As for *parallel* networks, these networks process all tasks simultaneously. Its final representation motif can be estimated by collapsing the final representation motifs in the main diagonal of the *task-switching* network (or of the *independent* network, as they are indistinguishable). Mathematically, we can collapse the task-specific blocks in two ways, by computing the normalized *union* of the different tasks or by computing their *intersection*:

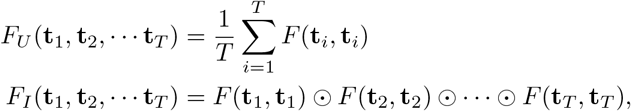

where ⊙ represents the Hadamard (or element-wise) product. We then assumed that the correlation matrix **R** converges to *F*_*p*_ = *cF*_*U*_ + (1 − *c*)*F*_*I*_ in the final representation (i.e., that of the last hidden layer), with the parameter *c* ∈ [0, 1].

S1 Fig S20 compares the empirical final block motif of the RDM in a 10-layer *parallel* network with its prediction for *c* = 0.5. The coefficient of determination *R*^2^ in all conditions was *>* 0.95. We also adjusted the parameter *c* for each case using least mean squares algorithm, however, adjusted *c* values showed no clear trend with the increased number of tasks. Furthermore, the difference in the residuals (and consequently *R*^2^) between *c* = 0.5 and adjusted *c* was barely noticeable.

The RDM of *parallel* networks is composed by tiling the same multitask block. Hence, the multitask tile representation MR_tile_ is a built-in block to compute complete, task and shared representations in *parallel* networks:

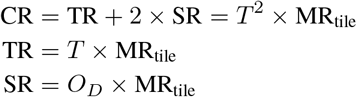

The multitask tile representation can be estimated by averaging *F*_*p*_:

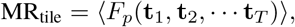

S1 Fig S20 also compares the predicted mean similarity representation with that in the empirical final block motif of the RDM, showing a good estimation of the multitask tile representation. Remarkably, the larger the number of processed tasks in *parallel* networks, the lower the task and shared representations become.

### Multidimensional scaling

We applied multidimensional scaling (MDS) (Kruskal, 1964) to the RDM of each hidden layer (Jazayeri & Ostojic, 2021; Bernardi et al., 2020). For this, the RDM is taken as a distance matrix between each *i* and *j* position. We project this high-dimensional distribution of distances into a two-dimensional embedding using the MDS implementation in *scikit-learn* (Pedregosa et al., 2011). The maximum number of iterations was set to 1000 to guarantee convergence, with a tolerance of 10^−5^. We compare the results of the MDS analysis in three conditions: context only in the first layer, context in all layers and context only in the last layer (S1 Fig S6). In addition, the MDS analysis was also applied to project the task similarity matrix to 2D (S1 Fig S13).

### Linear decoder

We used a generalized linear model to decode the information processed in each layer and its evolution as processing progressed through the feedforward network architecture (S1 Fig S14) (Alain & Bengio, 2016; Kriegeskorte & Douglas, 2019). We applied the *scikit-learn* implementation of the multiclass logistic regression model (Pedregosa et al., 2011). The maximum number of iterations was set to 8000 to guarantee convergence with a tolerance of 10^−3^. We used L2-norm regularization and the cross-entropy function for the loss.

The linear decoder was applied to the activation matrix of each layer, assembled as described in the Section Representational dissimilarity matrix. We first divided the data into a train set and test set following a 90/10 split and used digit labels (0–9) to identify the corresponding task label (see Table 1) and generalized congruency (see Section Generalized congruency). The results from the analysis were averaged across 10 realizations with different random seeds (S1 Fig S14).

### Parameter scalability with the number of tasks

We can estimate the total number of parameters for *independent, parallel* and *task-switching* networks (respectively, *P*_s_, *P*_p_ and *P*_ts_). For a fixed number of hidden neurons *H*, hidden layers *L*, input size *I* and output size *O*, the number of parameters grows linearly with the number of tasks in 𝕋. Assuming there are bias parameters in all layers, the number of parameters as a function of the number of tasks *P* (*T*) in *independent* and *parallel* networks is:

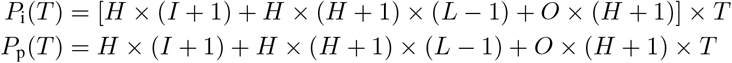

Note that in our models, neurons in the *task-switching* network have as many contexts biases as tasks are in T, unlike neurons in the *independent* and *parallel* networks that have only a single bias parameter. However, the number of parameters in the *task-switching* network can be further restricted keeping context biases the same (*S*) for neurons belonging to a same layer, compared to the different condition (*D*) where context biases are all different:

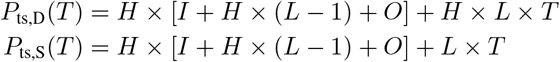

Considering the *S* condition is based on biological foundations, assuming low variability in the context input from PFC to downstream neurons within a brain area (e.g., neurons in the cortical visual area V4), in comparison to significantly larger variability across brain regions (e.g., neurons in visual area V1 vs. V4).

The dependency with *T* is linear in the three network architectures above, but the slope in the *task-switching* network *S* condition is reduced from *H* × *L*, to *L*. In terms of scalability, the number of parameters of *task-switching* networks in the *S* condition will grow much slower than that of *parallel* networks, since *O*× (*H* + 1) ≫*L*. This is because the number of neurons within a layer *H* is typically much larger than the number of layers *L* in a network. In fact, S1 Figure S16 displays the number of parameters in the different architectures for up to 100 tasks. We we can see that the line for the *task-switching* networks appears flat for the *S* condition, in contrast to other cases.

### Clustering of neural activations

Similarly as in the assembling of the RDM, we averaged the neural activity for the last layer for each digit, creating a matrix of size *S*×*H*, where *S* is the number of stimulus types and *H* is the number of neurons in the hidden layer. We gathered the averages of activity for the different tasks to construct a matrix of size *S*×*T*×*H*, where *T* are the number of tasks. Each independent column conforms the average of activations across tasks and digits for a single hidden neuron. We normalized the activity by the maximum value so the new maximum is set to 1. Upon these activations, we applied K-means clustering (Lloyd, 1982; Pedregosa et al., 2011).

The analysis associated each hidden neuron to a cluster and we then calculated the average activity of each cluster per task and digit (see S1 Fig S22). We ran K-means for different number of clusters and inspected their average activation patterns. We found that the best number of clusters corresponds to *k* = 3, since three clusters distinguish neurons contributing to either output response of each task as well as silent neurons.

For *k <* 3, the specificity for the output response was first lost (*k* = 2), and then also the specificity for sparsity (defined as the proportion of silent neurons), when a single cluster was considered. The proportion of silent neurons in the last hidden layers was large (see S1 Fig S21), which explained why K-means separated first silent from non-silent neurons. Finally, for *k >* 3, K-means separated clusters by activation strength rather than by activation pattern (see S1 Fig S22).

## Code availability

The code for replicating the network models and the results of the analyses presented in this manuscript is available in this GitHub repository.

## SI Supplementary Figures

**S1 Fig S1:**
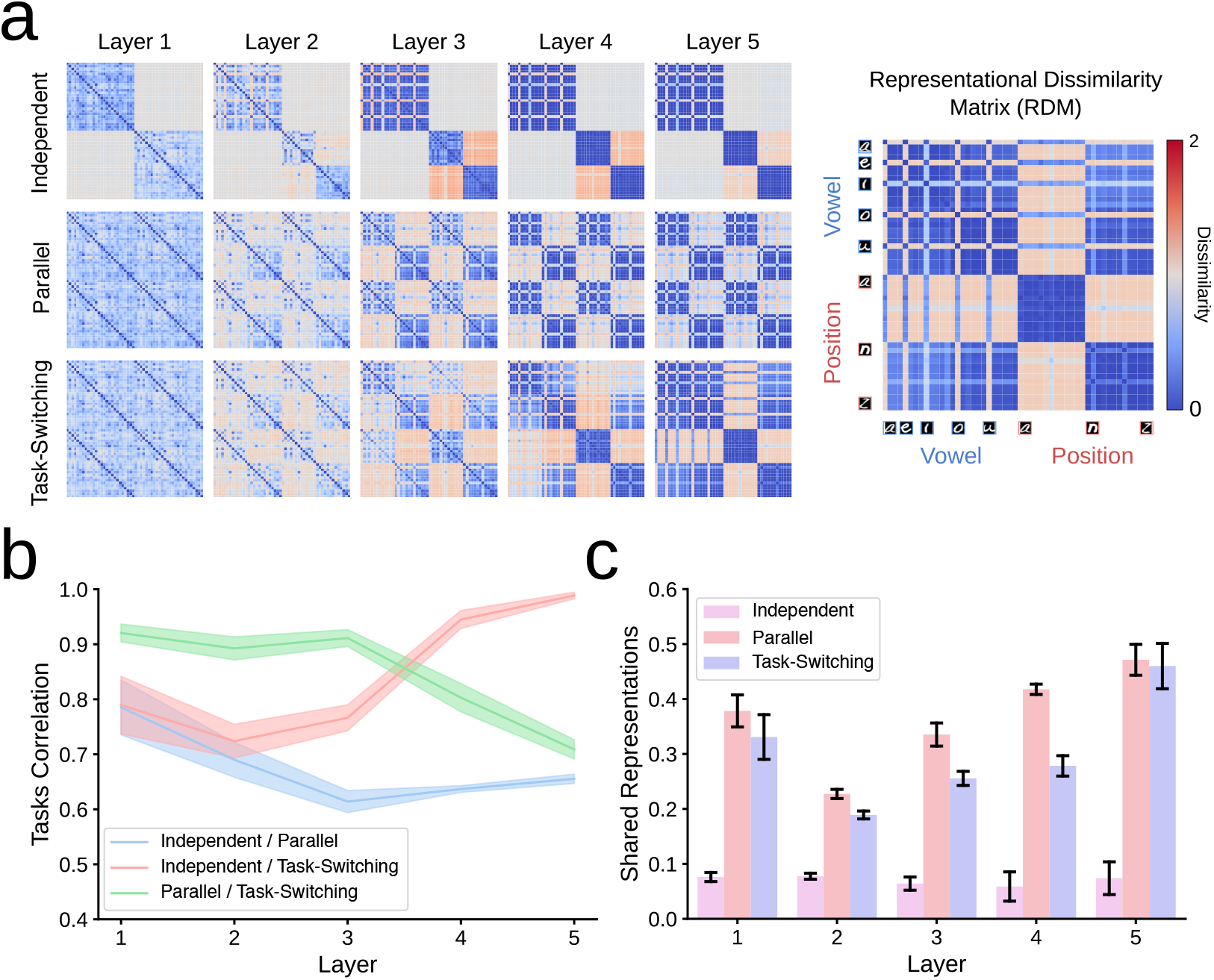
Neural representations for *independent, parallel* and *task-switching* networks for two distinct two-alternative choice tasks based on letter processing: {*vowel, position*}. (a) Average representational dissimilarity matrix (RDM) for the three architectures across 10 different initializations (*runs*). (b) Task correlation between models measured by the Pearson correlation between RDMs’ main diagonal blocks (Mean and SD over 10 different runs). (c) Shared representations between different tasks in each model measured by the sum of absolute correlation values in RDMs’ off-diagonal blocks (Mean and SD over 10 runs).

**S1 Fig S2:**
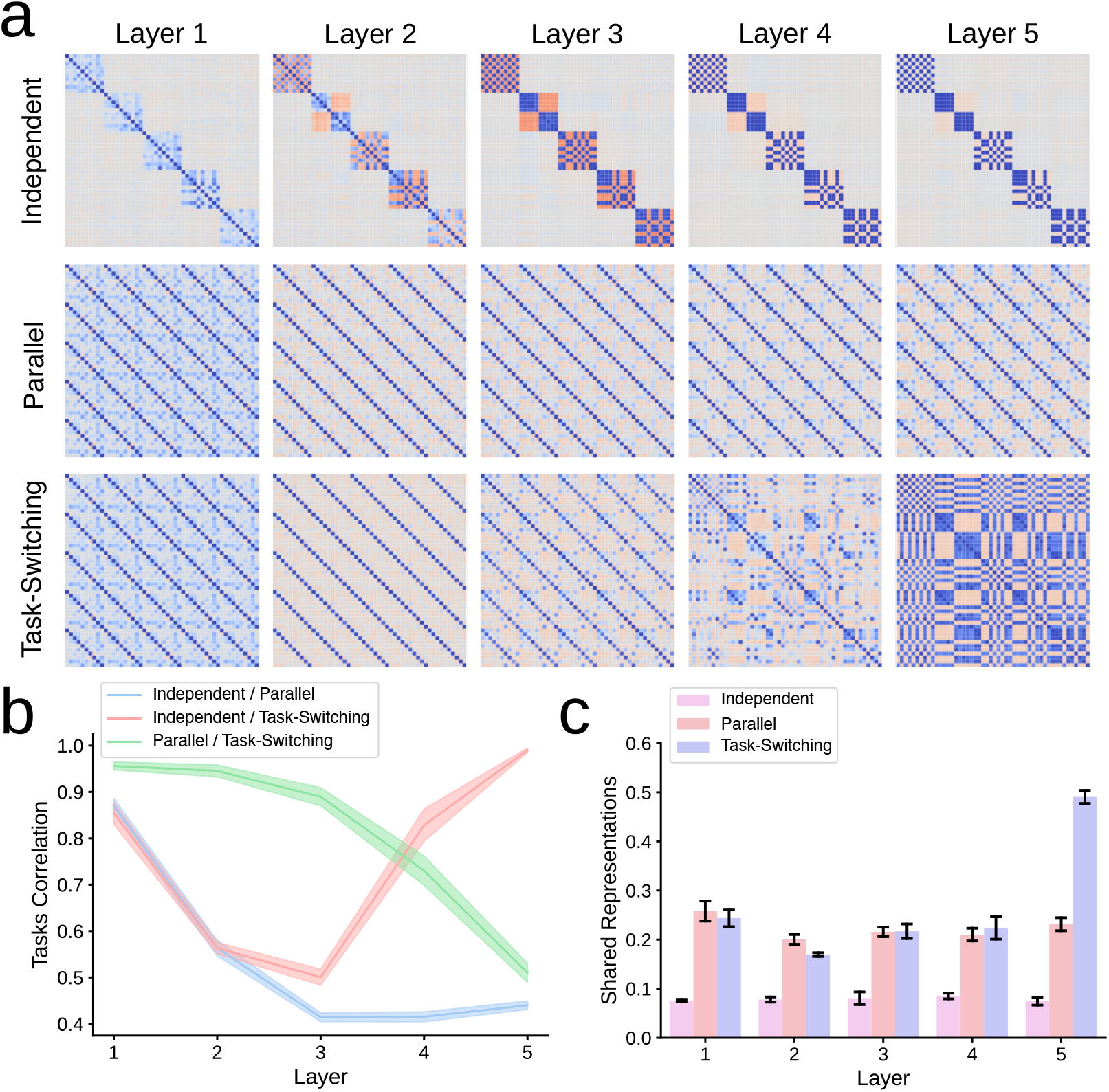
Neural representations for *independent, parallel* and *task-switching* networks for five distinct two-alternative choice tasks based on digit processing: {*parity, value, prime, fibonacci, multiple of 3*}. (a) Average representational dissimilarity matrix (RDM) for the three architectures across 10 different initializations (*runs*). (b) Task correlation between models measured by the Pearson correlation between RDMs’ main diagonal blocks (Mean and SD over 10 different runs). (c) Shared representations between different tasks in each model measured by the sum of absolute correlation values in RDMs’ off-diagonal blocks (Mean and SD over 10 runs).

**S1 Fig S3:**
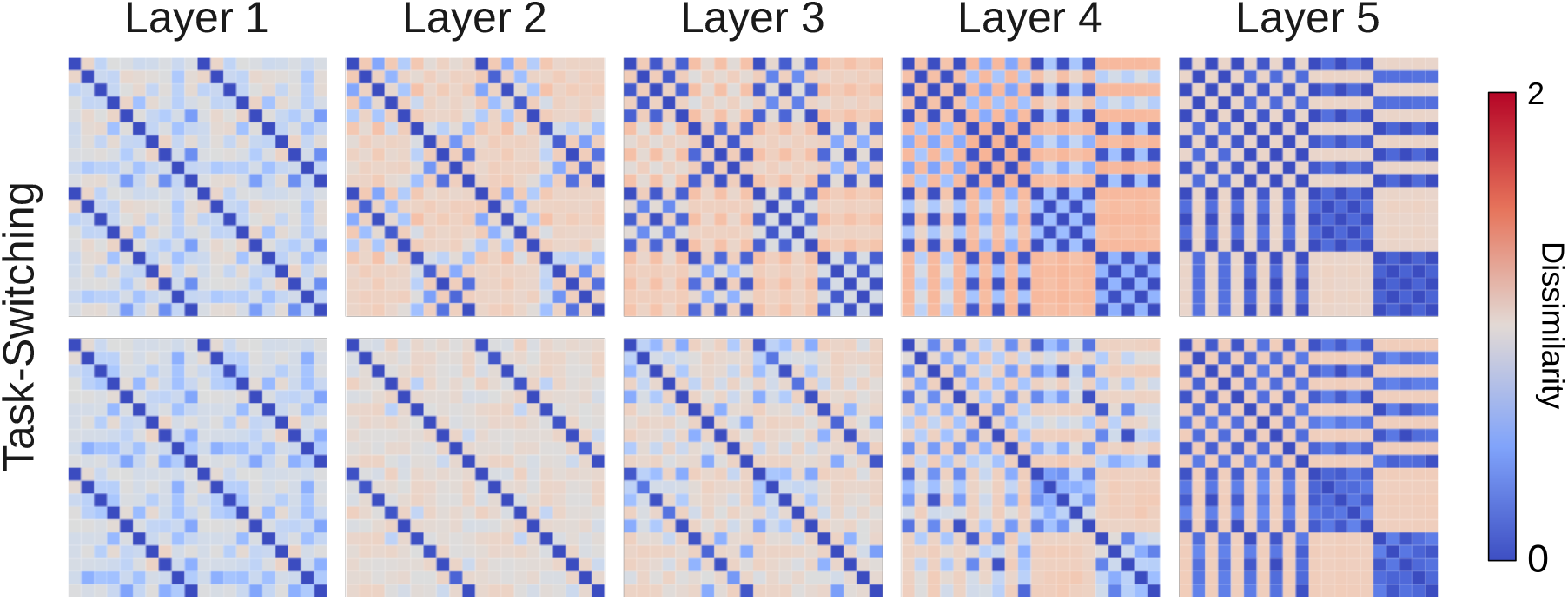
RDMs for five-layer *task-switching* networks for 2 tasks, {*parity, value*} (top) and 5 tasks, {*parity, value, prime, fibonacci* and *multiples of 3*} (bottom). We cropped the RDM for 5 tasks to better compare *parity* and *value* in the two cases.

**S1 Fig S4:**
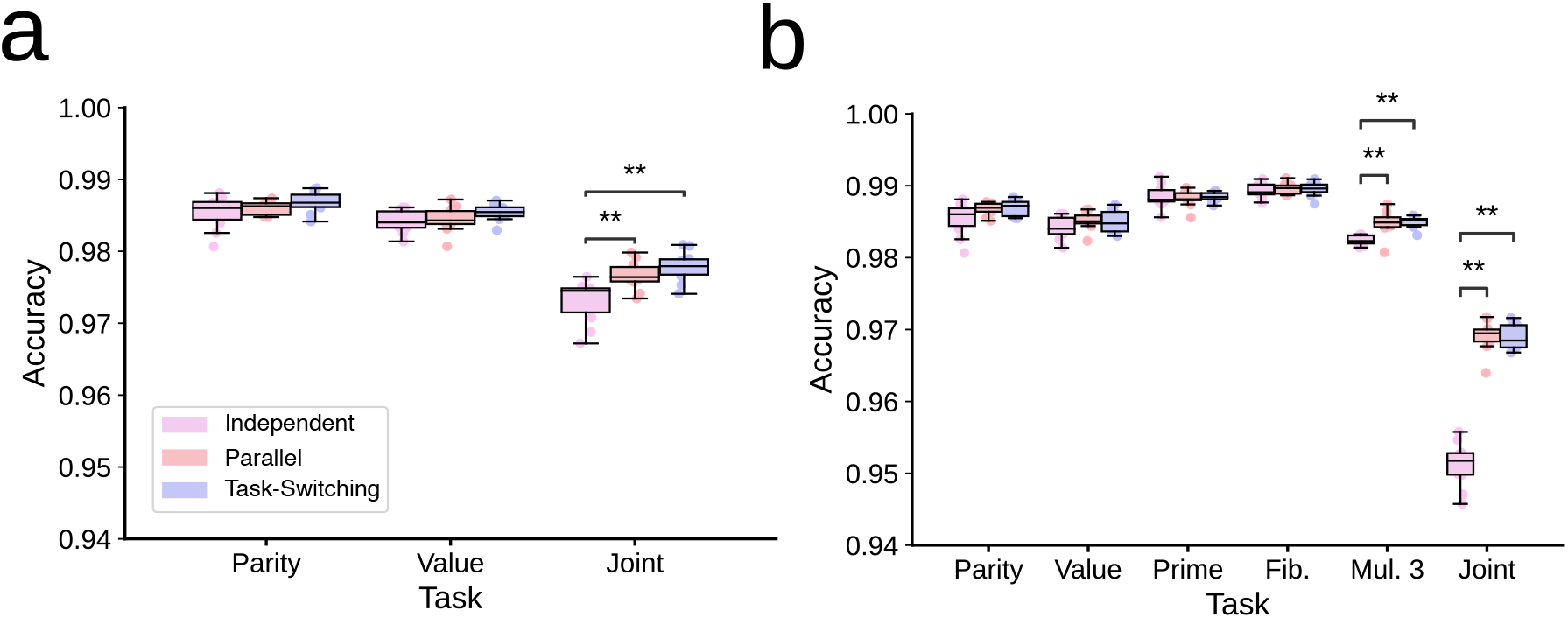
Accuracy for five-layer *independent, parallel* and *task-switching* networks. **(Wilcoxon Signed-Rank Test**, *p <* 0.001**) for {*parity, value*} (a), and for {*parity, value, fibonacci, prime, multiple of 3*} (b)**.

**S1 Fig S5:**
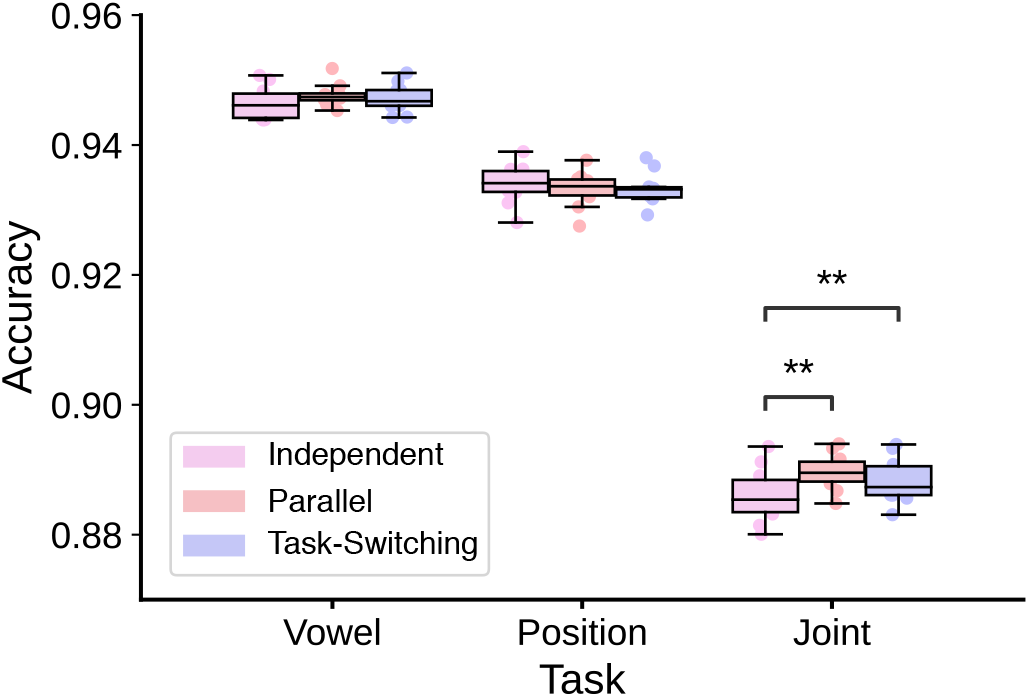
Accuracy for five-layer *independent, parallel* and *task-switching* networks for {*letter, position*}. ** denotes *p <* 0.001 in the Wilcoxon Signed-Rank Test.

**S1 Fig S6:**
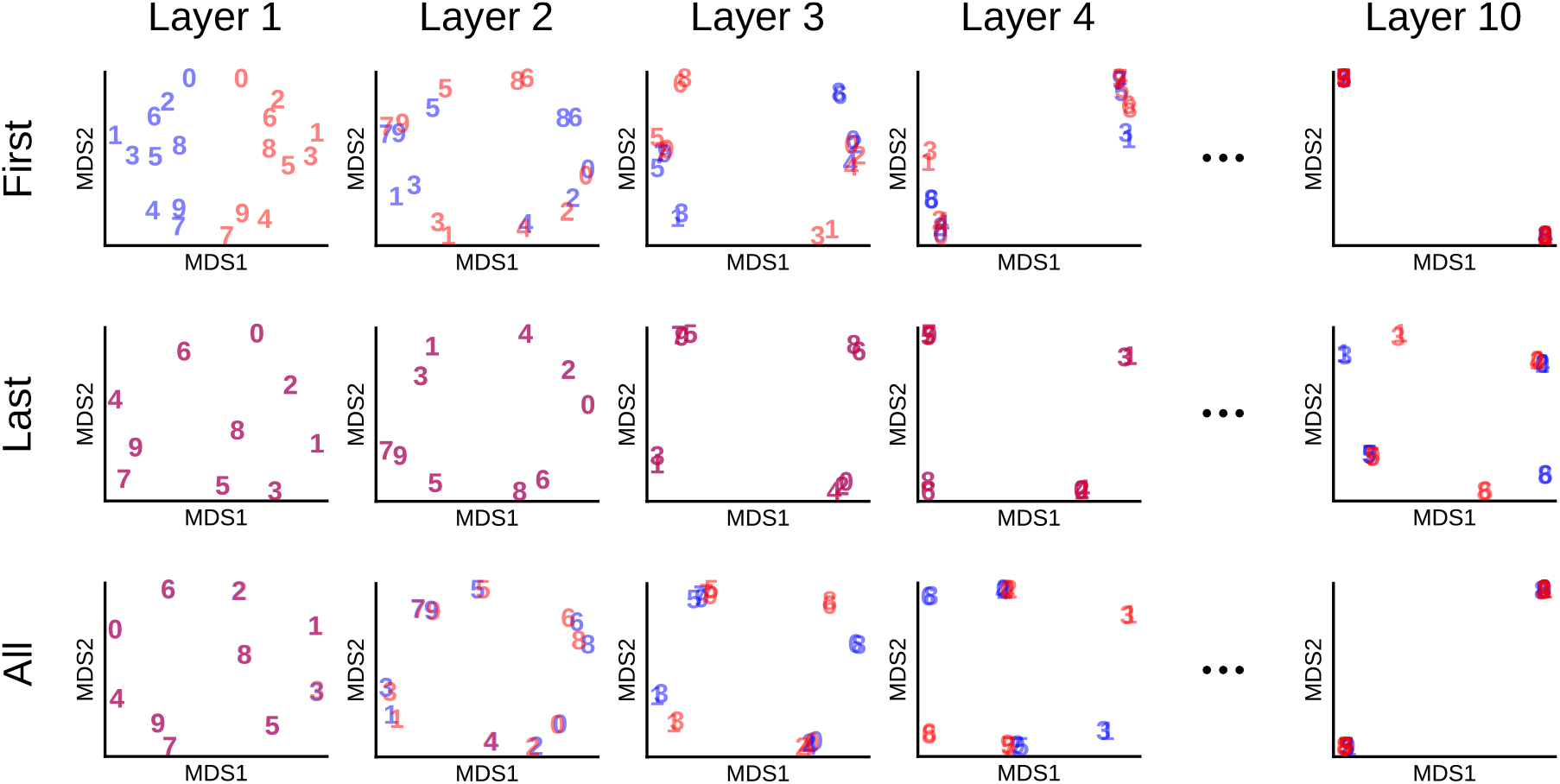
Multidimensional scaling of RDMs obtained for *task-switching* networks when context is added only in the first layer (First), last layer (Last) or in all layers (All) for {*parity, value*} *task-switching* schedules. Blue and red digits denote low-dimensional representations of task *parity* and *value*, respectively.

**S1 Fig S7:**
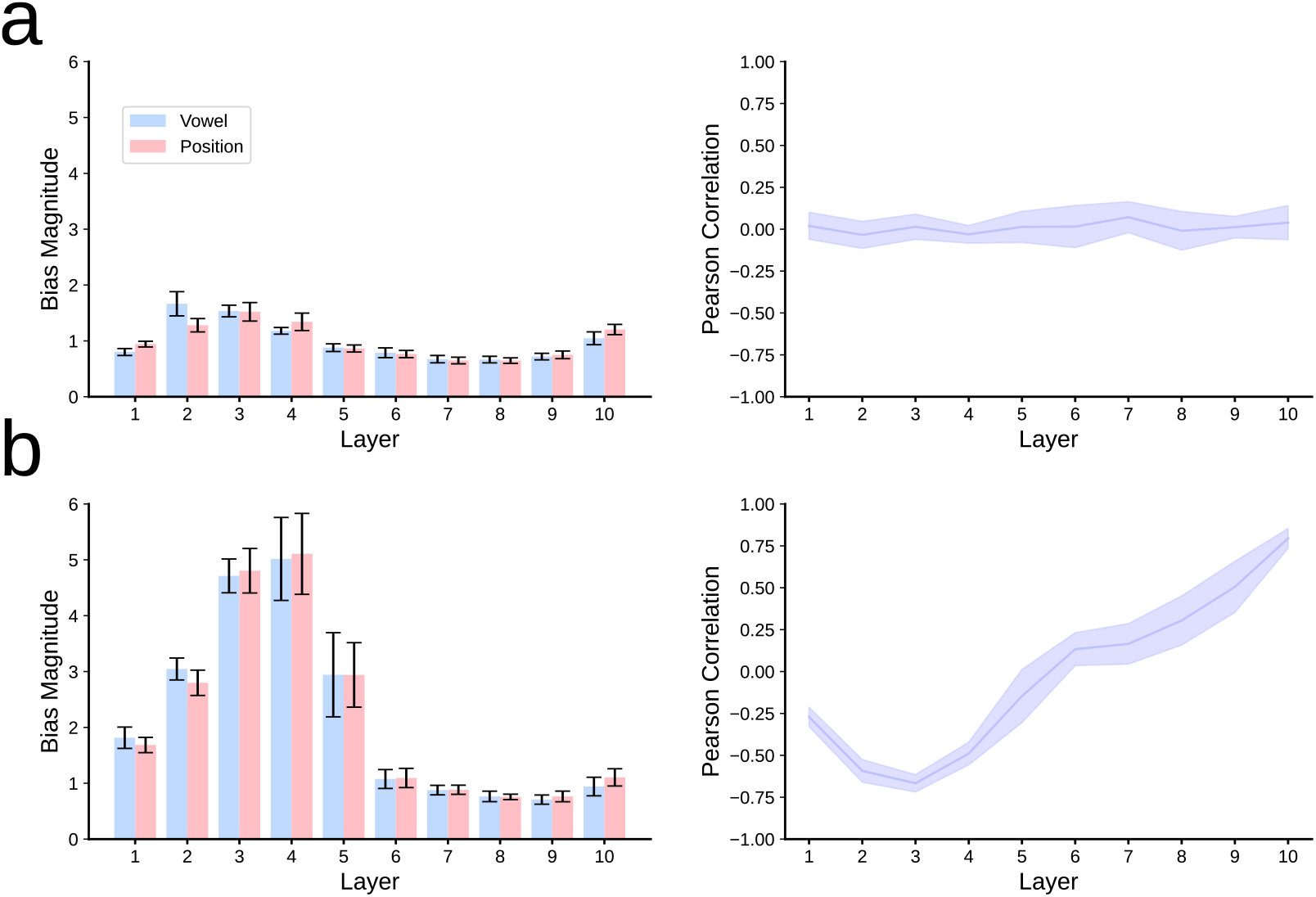
Bias connection strength and its correlation in *independent* networks under letter processing. Bias magnitude (a,c) and Pearson correlation (b,d) between input biases in *independent* networks (a,b), compared to *task-switching* networks (c,d) for the {vowel, position} tasks (Mean and SD across 10 runs). Note that input biases are associated with context inputs only in *task-switching* networks. The magnitude of the bias is calculated using the *L*2-norm (see Section Context bias strength).

**S1 Fig S8:**
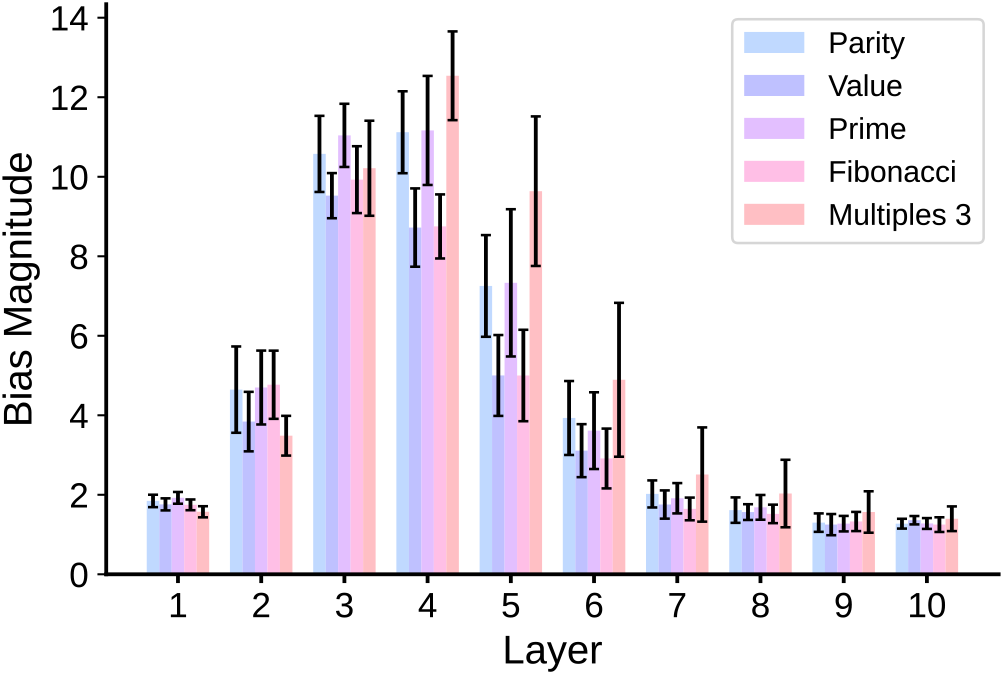
Bias connection strength in *independent* networks trained in 5 digit tasks, {*parity, value, prime, fibonacci* and *multiples of 3*}. The magnitude of the bias is calculated using the L2-norm (see Section Context bias strength). Mean and SD is shown across 10 runs.

**S1 Fig S9:**
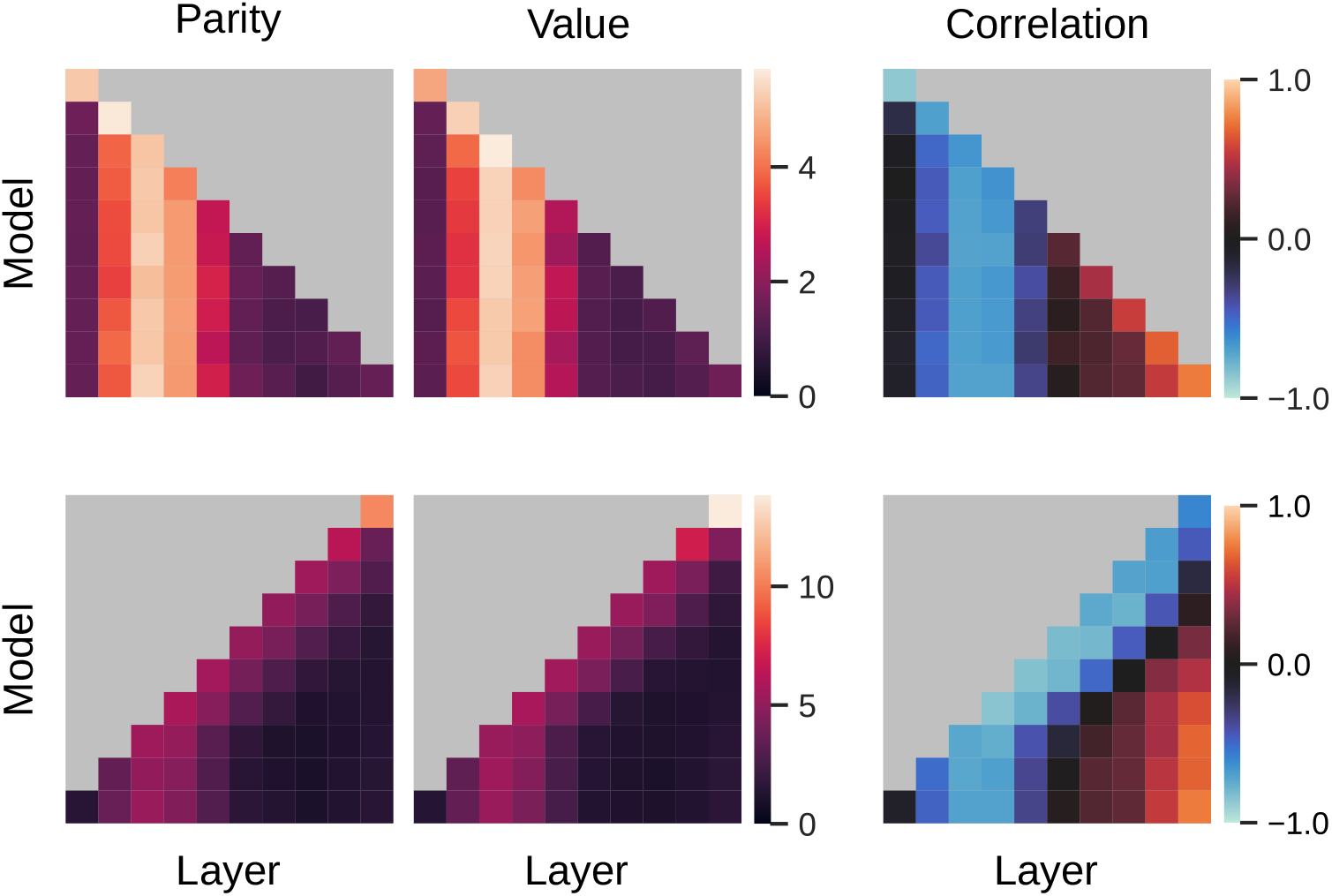
Mean context connection strength and correlation between contexts (multiple contexts). (a) Contexts added from layer 1 to layer 10. (b) Contexts added from layer 10 to layer 1. Networks of 10 layers for 10 different runs. The gray background denotes the presence of a layer without context.

**S1 Fig S10:**
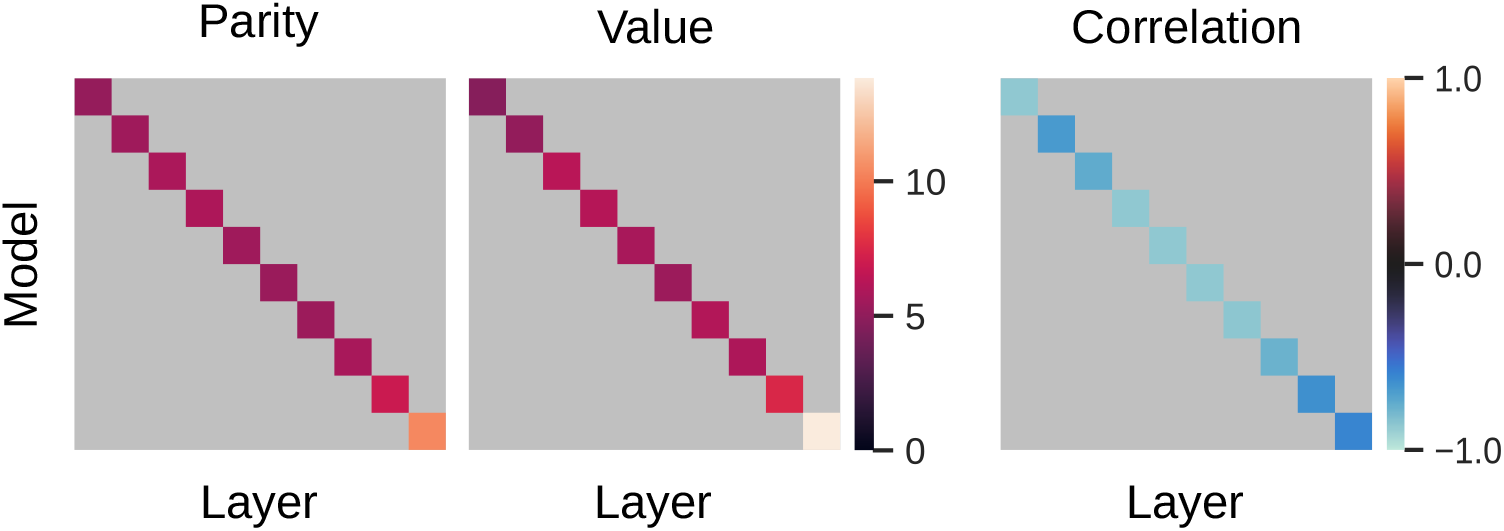
Mean context connection strength and correlation between contexts (single context). Networks of 10 layers for 10 different runs. The gray background denotes the presence of a layer without context.

**S1 Fig S11:**
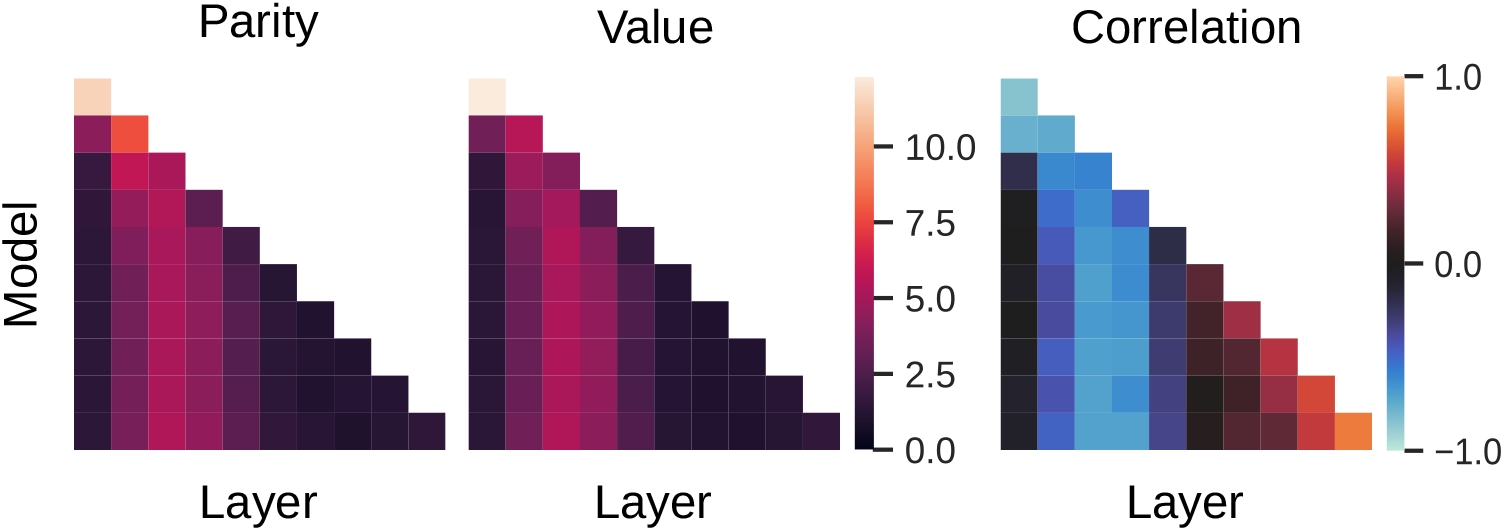
Mean context connection strength and correlation between contexts (single context (multiple architectures). Networks have different number of layers, from 1 to 10. Contexts are present in all layers. The white background denotes the absence of a layer.

**S1 Fig S12:**
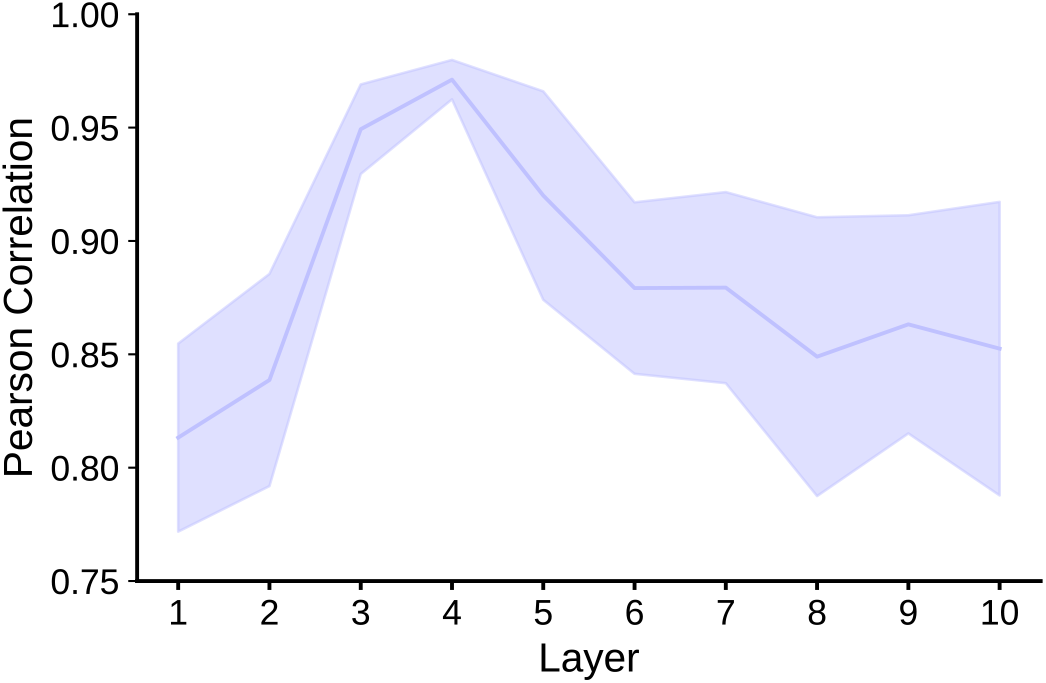
Correlation between task similarity and congruency. The task-switching network was trained in five digit tasks: {*parity, value, prime, fibonacci* and *multiples of 3*}. Mean and SD is shown across 10 runs.

**S1 Fig S13:**
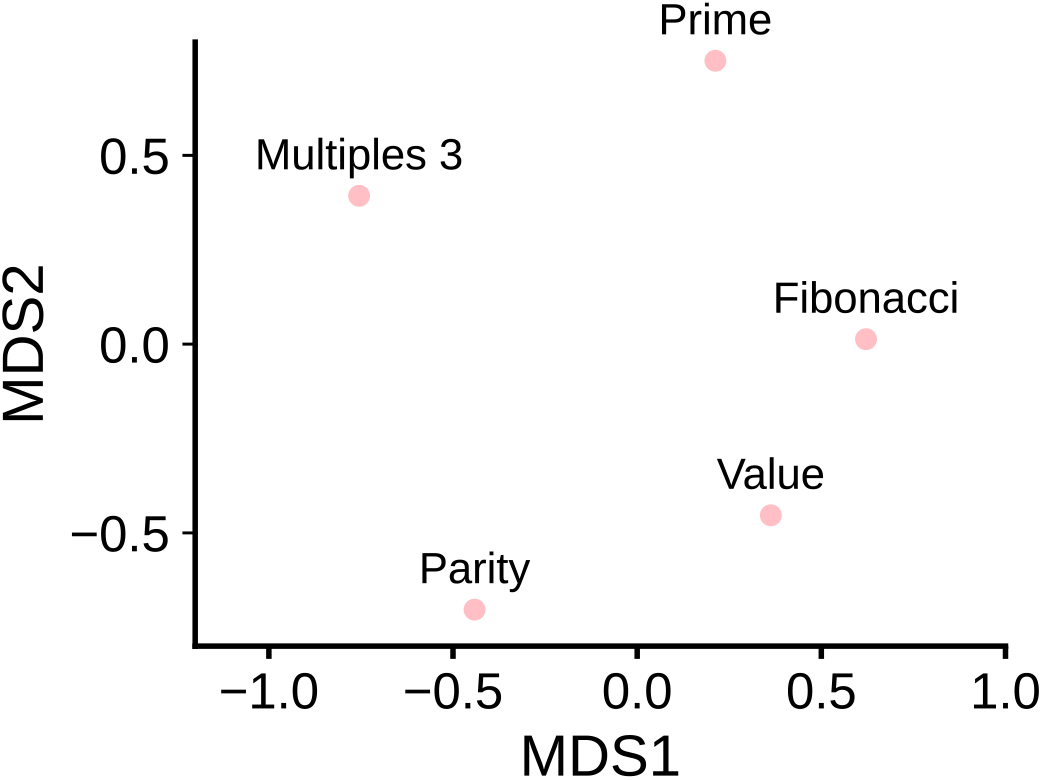
2D embedding of the task similarity matrix for *task-switching* networks trained for 5 binary digit tasks, {*parity, value, prime, fibonacci* and *multiples of 3*}.

**S1 Fig S14:**
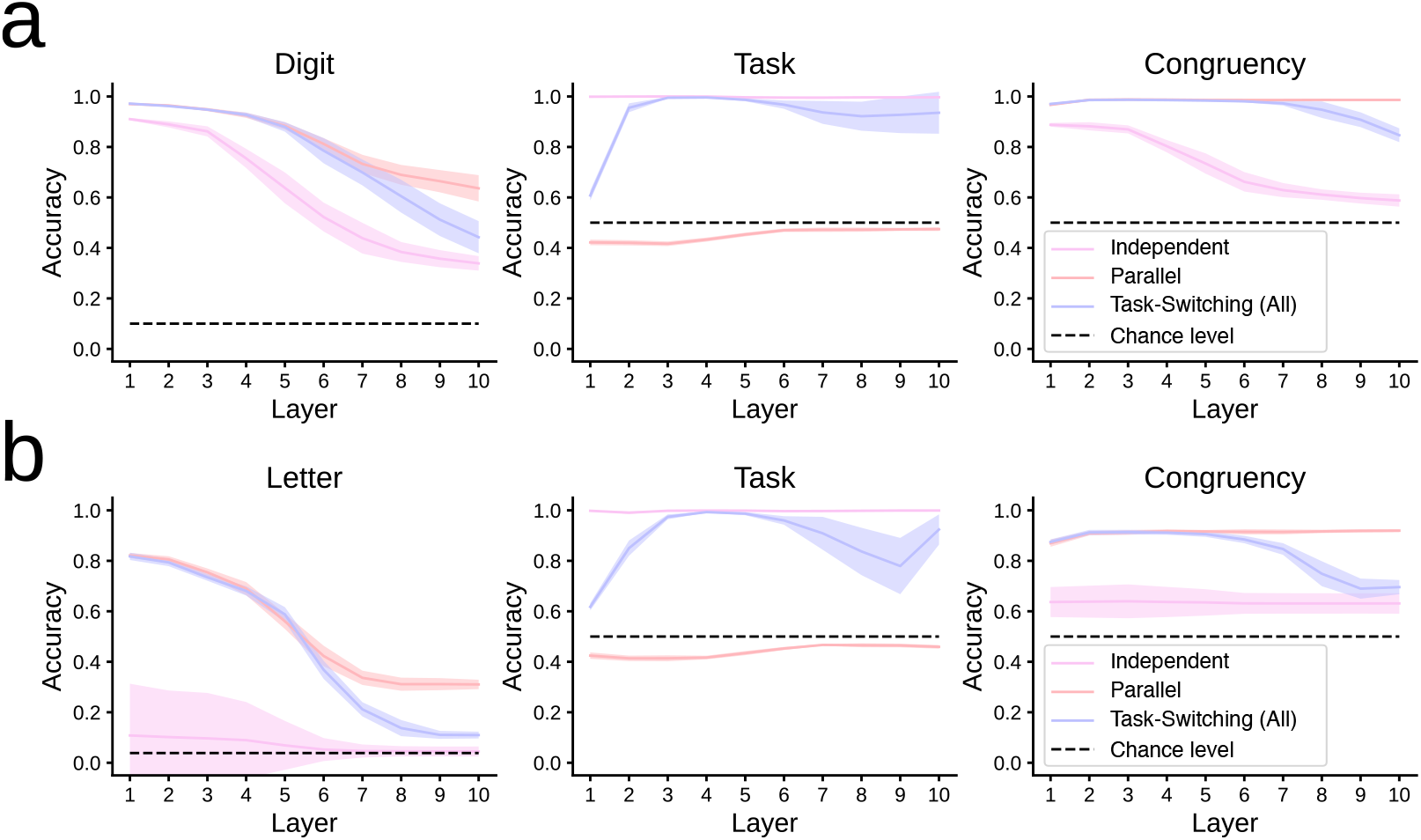
Linear decoders of neural activity for input type (0–9 for digits and a–z for letters), task output (0/1 labels) and generalized congruency (yes/no). (a) Decoders for *independent, parallel* and *task-switching* with context in all layers for {*parity, value*}. (b) Decoders for *independent, parallel* and *task-switching* with context in all layers for {*vowel, position*}. Mean and SD values are shown across 10 runs.

**S1 Fig S15:**
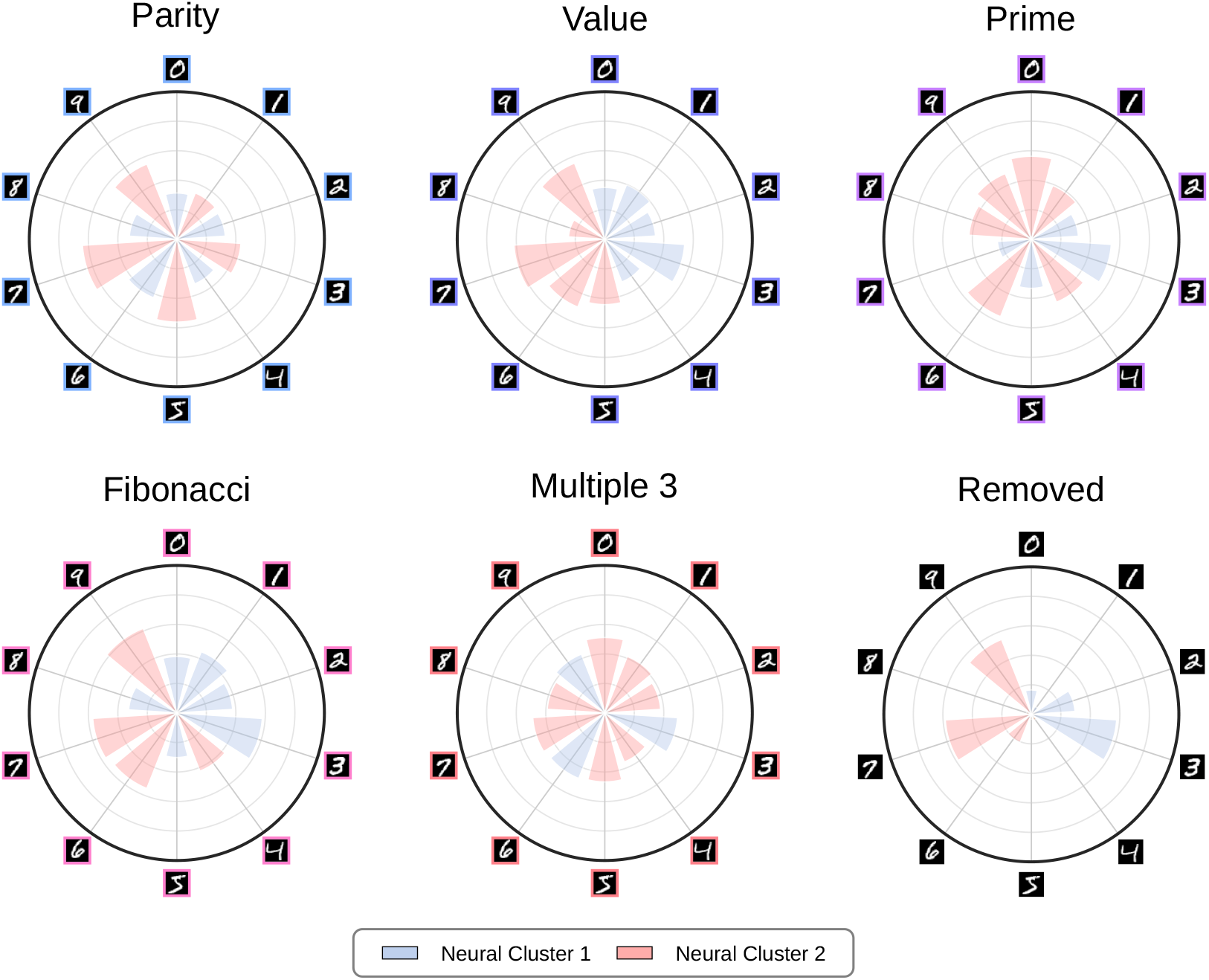
Neural clustering for the {*parity, value, prime, fibonacci, multiple of 3*} *task-switching* schedule.

**S1 Fig S16:**
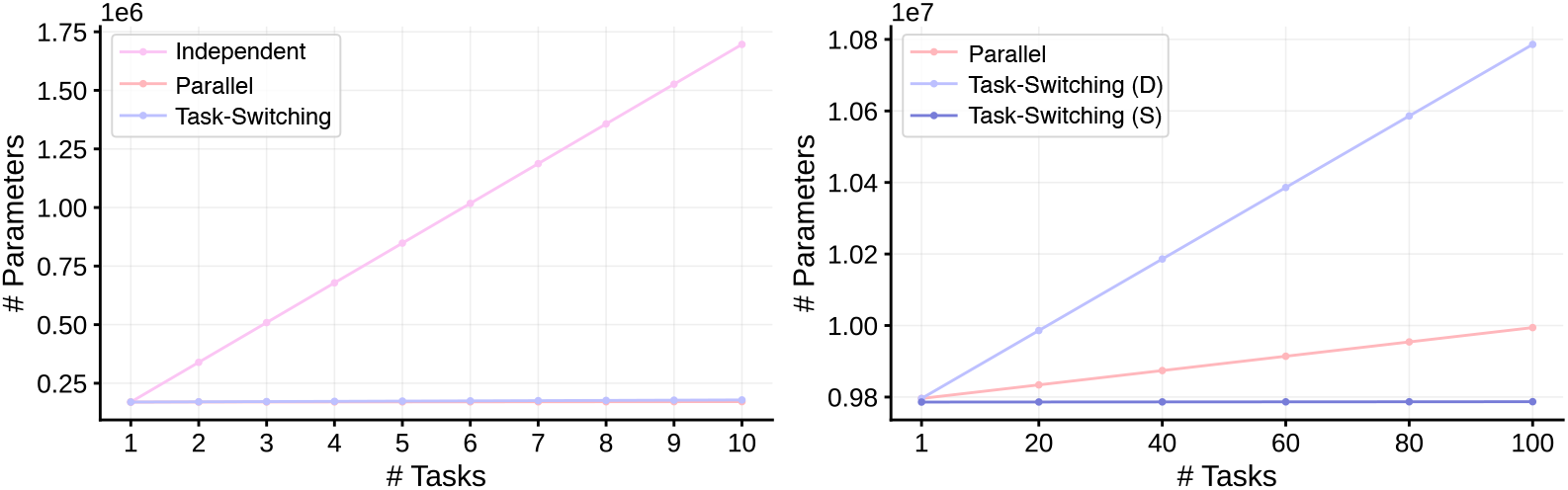
Parameters in neural networks used in *task switching* for two-alternative choice tasks. All three models grow linearly with the number of tasks. The number of parameters for the *independent* networ grows much faster than for *parallel* and *task-switching* networks (left). *Parallel* networks lie in between of two flavors of *task-switching* networks: (D) denotes different context parameters in all layers, whereas (S) denotes same contexts for all neurons within a layer. We assumed a fixed number of tasks (10 in the left panel, 100 in the right panel), for a fixed number of hidden layers (10), neurons in each layer (100), and using MNIST digit images as input (28 × 28 pixels).

**S1 Fig S17:**
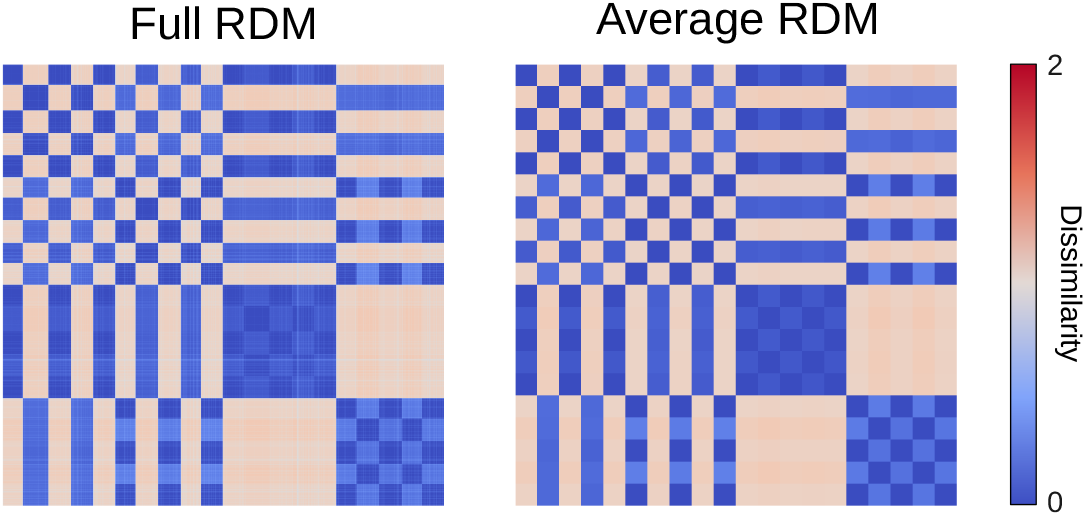
Full vs. average RDMs. On the left, the RDM is formed by the product of neural activations in tasks (in this case 2) across the total number of test images (10,000). We have sorted the elements in the test set so that they are grouped together by digit. On the right, the RDM is formed after averaging the activity across the different instances of the same digit.

**S1 Fig S18:**
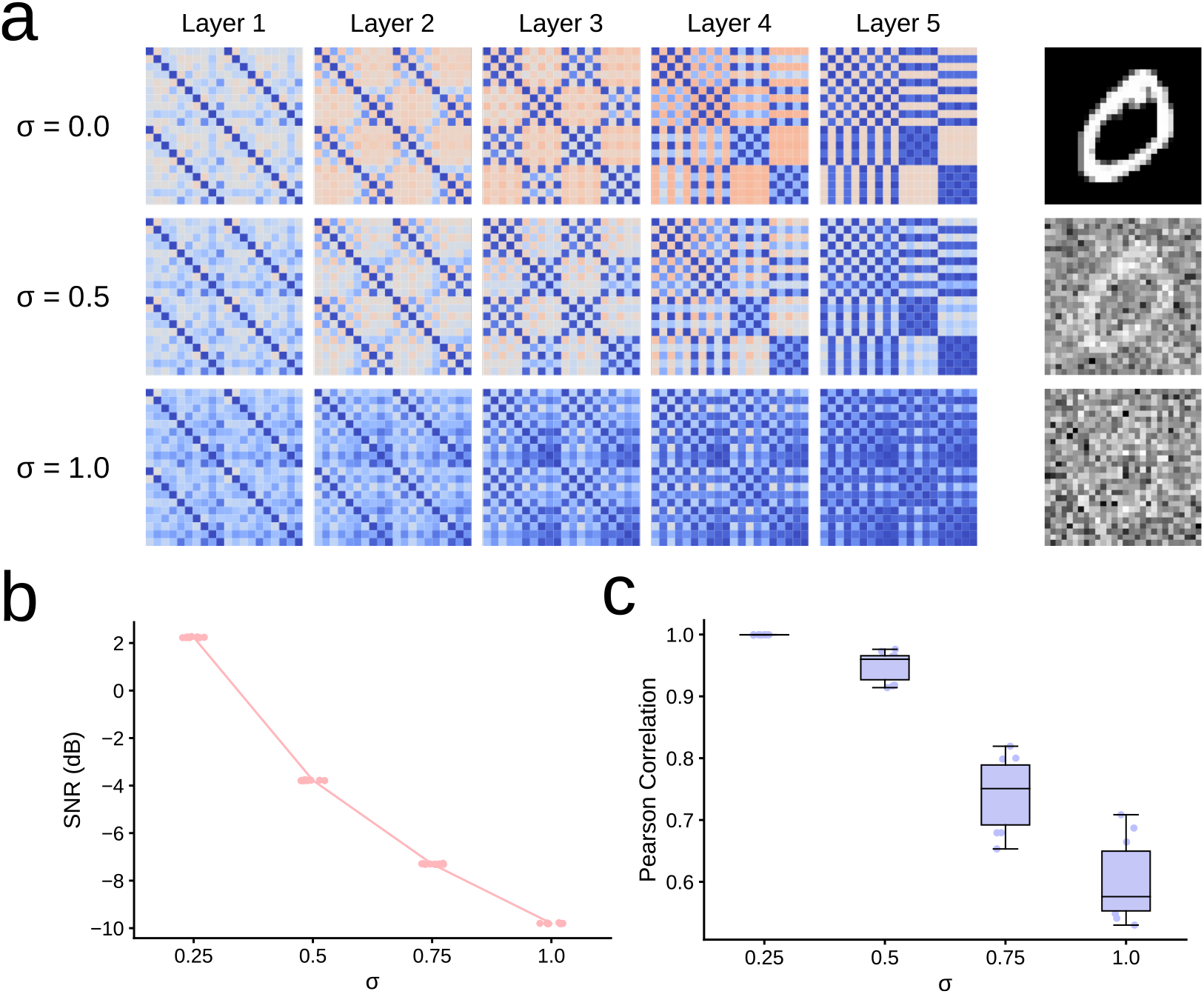
Robustness of RDMs against noise. a) We have added Gaussian noise with standard deviation *σ* to the test images and constructed their RDMs. RDMs are altered significantly only when the noise strongly masks the digit. b) Signal-to-noise ratio for different values of *σ*, defined in dB as 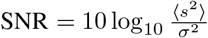, where *s* denotes the pixels’ intensity. c) Correlation of noisy RDMs with the RDM without noise. Even when the power of noise is considerable, the correlation with the noiseless RDM is high.

**S1 Fig S19:**
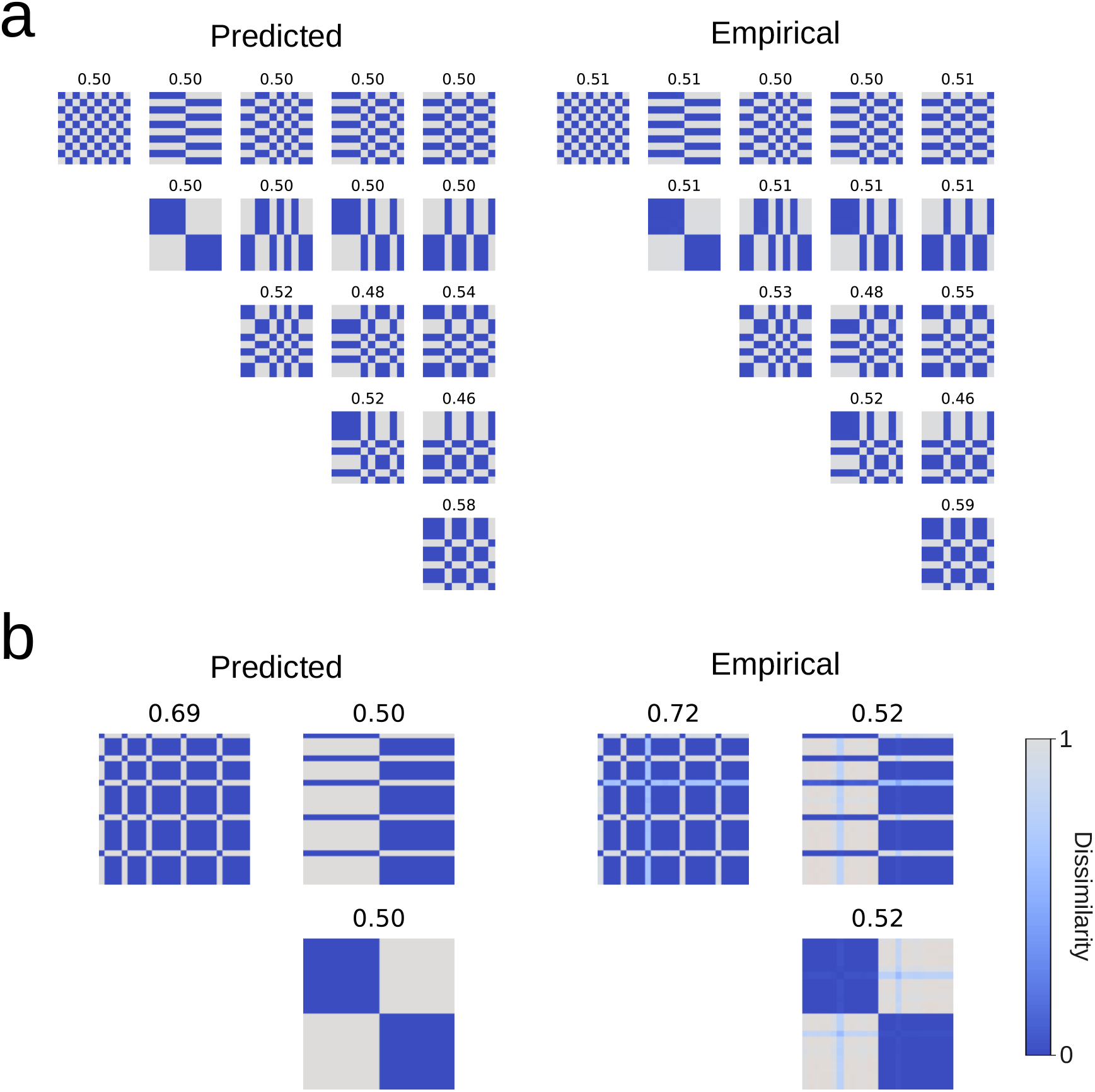
Predicted and empirical final representation motifs in the last of 10 layers of the *task-switching* network. a) for {*parity, value, prime, fibonacci* and *multiples of 3*} digit tasks. b) for {*vowel* and *position*} letter tasks. The title of each block denotes the averaged similarity representation.

**S1 Fig S20:**
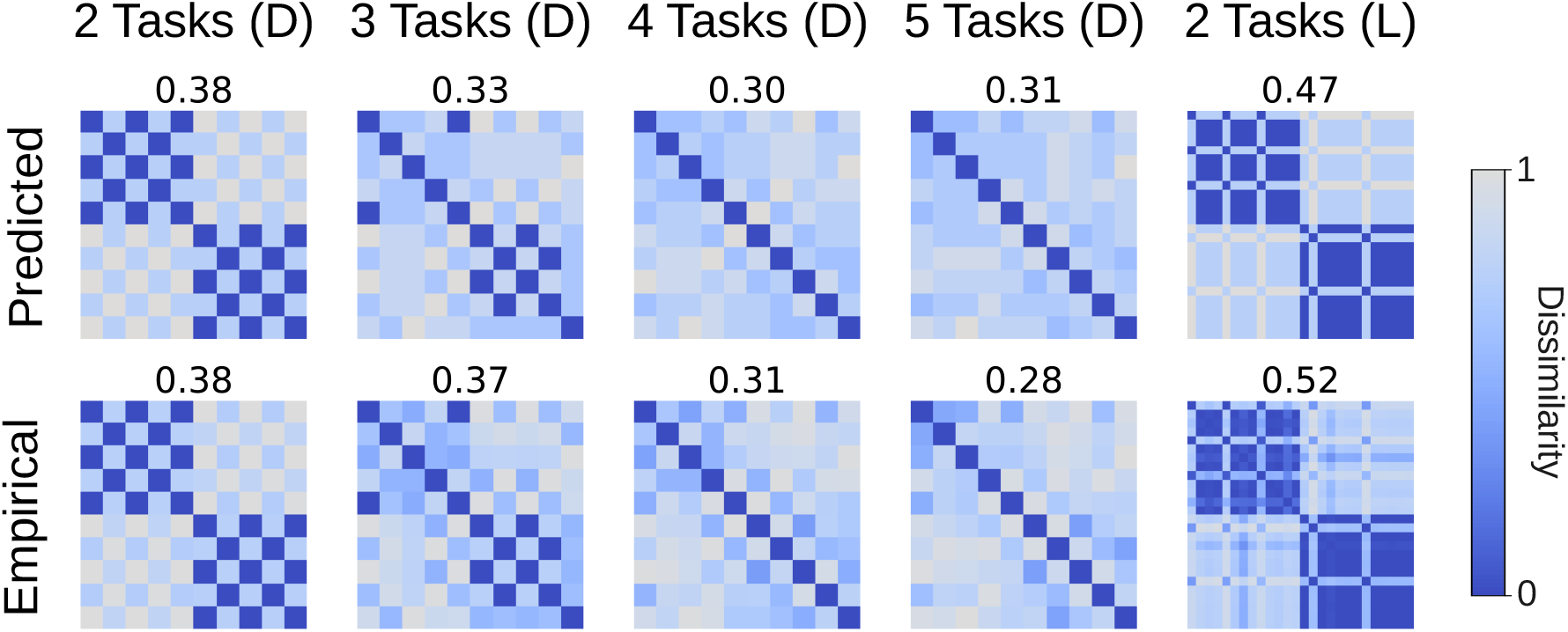
Predicted and empirical final representation motifs in the last of 10 layers of the *parallel* network. First columns show two-alternative choice tasks with digits (D): {*parity, value*}, {*parity, value, prime*}, {*parity, value, prime, fibonacci*}, and {*parity, value, prime, fibonacci, multiples of 3*}. The last column shows a two-alternative choice tasks with letters (L), {*vowel, position*}. The title of each block denotes the averaged similarity representation.

**S1 Fig S21:**
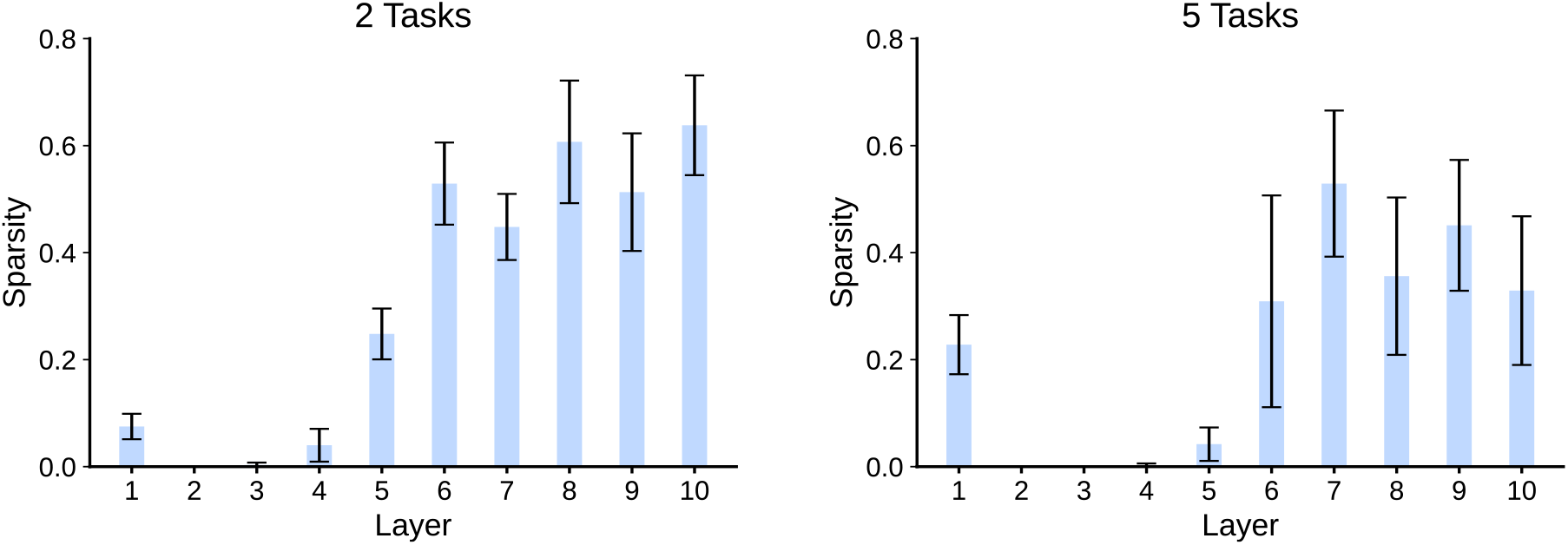
Sparsity for task-switching networks. Networks were trained with {*parity, value*} and {*parity, value, prime, fibonacci* and *multiples of 3*}, respectively. Mean sparsity and SD shown across 10 runs.

**S1 Fig S22:**
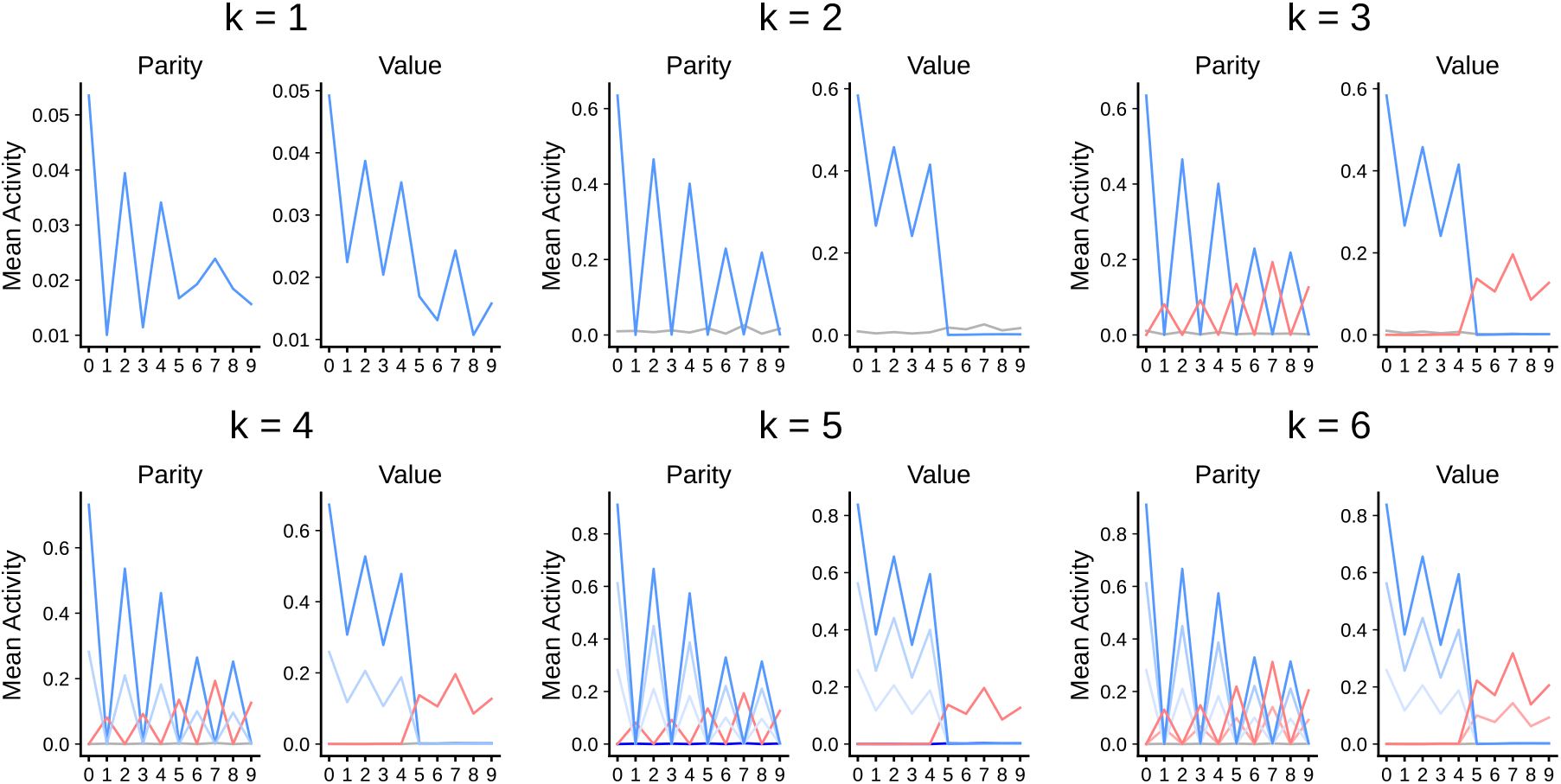
Activity patterns of the last hidden layer. Resulting activity patterns when imposing *k* clusters.

## References

Alain, G., & Bengio, Y. (2016). Understanding intermediate layers using linear classifier probes. arXiv preprint 1610.01644.

Ardid, S., Vinck, M., Kaping, D., Marquez, S., Everling, S., & Womelsdorf, T. (2015). Mapping of functionally characterized cell classes onto canonical circuit operations in primate prefrontal cortex. Journal of Neuroscience, 35(7), 2975–2991.

Ardid, S., & Wang, X.-J. (2013). A tweaking principle for executive control: neuronal circuit mechanism for rule-based task switching and conflict resolution. Journal of Neuroscience, 33(50), 19504–19517.

Bengio, Y., Courville, A., & Vincent, P. (2013). Representation learning: A review and new perspectives. IEEE transactions on pattern analysis and machine intelligence, 35(8), 1798–1828.

Bernardi, S., Benna, M. K., Rigotti, M., Munuera, J., Fusi, S., & Salzman, C. D. (2020). The geometry of abstraction in the hippocampus and prefrontal cortex. Cell, 183(4), 954–967.

Botvinick, M. M., Braver, T. S., Barch, D. M., Carter, C. S., & Cohen, J. D. (2001). Conflict monitoring and cognitive control. Psychological review, 108(3), 624.

Brock, A., Donahue, J., & Simonyan, K. (2018). Large scale gan training for high fidelity natural image synthesis. arXiv preprint 1809.11096.

Caruana, R. (1997). Multitask learning. Machine learning, 28(1), 41–75.

Cohen, G., Afshar, S., Tapson, J., & Van Schaik, A. (2017). Emnist: Extending mnist to handwritten letters. In 2017 international joint conference on neural networks (ijcnn) (pp. 2921–2926).

Cohen, J. D., Dunbar, K., & McClelland, J. L. (1990). On the control of automatic processes: a parallel distributed processing account of the stroop effect. Psychological review, 97(3), 332.

Crawshaw, M. (2020). Multi-task learning with deep neural networks: A survey. arXiv preprint 2009.09796.

Demšar, J. (2006). Statistical comparisons of classifiers over multiple data sets. The Journal of Machine learning research, 7, 1–30.

Ebitz, R. B., & Hayden, B. Y. (2021). The population doctrine in cognitive neuroscience. Neuron, 109(19), 3055–3068.

Edelman, S. (1998). Representation is representation of similarities. Behavioral and brain sciences, 21(4), 449–467.

Egner, T., & Hirsch, J. (2005). Cognitive control mechanisms resolve conflict through cortical amplification of task-relevant information. Nature neuroscience, 8(12), 1784–1790.

Fischer, R., & Plessow, F. (2015). Efficient multitasking: Parallel versus serial processing of multiple tasks. Frontiers in psychology, 6, 1366.

Flesch, T., Balaguer, J., Dekker, R., Nili, H., & Summerfield, C. (2018). Comparing continual task learning in minds and machines. Proceedings of the National Academy of Sciences, 115(44), E10313–E10322.

Flesch, T., Juechems, K., Dumbalska, T., Saxe, A., & Summerfield, C. (2022). Orthogonal representations for robust context-dependent task performance in brains and neural networks. Neuron, 110(7), 1258–1270.

Flesch, T., Nagy, D. G., Saxe, A., & Summerfield, C. (2022). Modelling continual learning in humans with hebbian context gating and exponentially decaying task signals. arXiv preprint 2203.11560.

French, R. M. (1999). Catastrophic forgetting in connectionist networks. Trends in cognitive sciences, 3(4), 128–135.

Fusi, S., Miller, E. K., & Rigotti, M. (2016). Why neurons mix: high dimensionality for higher cognition. Current opinion in neurobiology, 37, 66–74.

Gilbert, C. D., & Sigman, M. (2007, Jun). Brain states: top-down influences in sensory processing. Neuron, 54(5), 677–696.

Glorot, X., Bordes, A., & Bengio, Y. (2011). Deep sparse rectifier neural networks. In Proceedings of the fourteenth international conference on artificial intelligence and statistics (pp. 315–323).

Goodfellow, I., Bengio, Y., Courville, A., & Bengio, Y. (2016). Deep learning (Vol. 1). MIT Press.

Grewal, K., Forest, J., Cohen, B. P., & Ahmad, S. (2021). Going beyond the point neuron: Active dendrites and sparse representations for continual learning. bioRxiv.

Hebb, D. O. (1949). The organization of behavior: A neuropsychological theory. New York: Wiley.

Ito, T., Klinger, T., Schultz, D., Murray, J., Cole, M., & Rigotti, M. (2022). Compositional generalization through abstract representations in human and artificial neural networks. Advances in Neural Information Processing Systems, 35, 32225–32239.

Ito, T., & Murray, J. D. (2021). Multi-task representations in human cortex transform along a sensory-to-motor hierarchy. bioRxiv.

Jazayeri, M., & Ostojic, S. (2021). Interpreting neural computations by examining intrinsic and embedding dimensionality of neural activity. Current opinion in neurobiology, 70, 113–120.

Johnston, K., Levin, H. M., Koval, M. J., & Everling, S. (2007). Top-down control-signal dynamics in anterior cingulate and prefrontal cortex neurons following task switching. Neuron, 53(3), 453–462.

Johnston, W. J., Palmer, S. E., & Freedman, D. J. (2020). Nonlinear mixed selectivity supports reliable neural computation. PLoS computational biology, 16(2), e1007544.

Kingma, D. P., & Ba, J. (2014). Adam: A method for stochastic optimization. arXiv preprint 1412.6980.

Kriegeskorte, N., & Douglas, P. K. (2019). Interpreting encoding and decoding models. Current opinion in neurobiology, 55, 167–179.

Kriegeskorte, N., & Kievit, R. A. (2013). Representational geometry: integrating cognition, computation, and the brain. Trends in cognitive sciences, 17(8), 401–412.

Kriegeskorte, N., Mur, M., & Bandettini, P. A. (2008). Representational similarity analysis-connecting the branches of systems neuroscience. Frontiers in systems neuroscience, 4.

Kruskal, J. B. (1964). Multidimensional scaling by optimizing goodness of fit to a nonmetric hypothesis. Psychometrika, 29(1), 1–27.

LeCun, Y., Bottou, L., Bengio, Y., & Haffner, P. (1998). Gradient-based learning applied to document recognition. Proceedings of the IEEE, 86(11), 2278–2324.

Li, H., Ouyang, W., & Wang, X. (2016). Multi-bias non-linear activation in deep neural networks. In International conference on machine learning (pp. 221–229).

Liu, S., Johns, E., & Davison, A. J. (2019). End-to-end multi-task learning with attention. In Proceedings of the ieee/cvf conference on computer vision and pattern recognition (pp. 1871–1880).

Lloyd, S. (1982). Least squares quantization in pcm. IEEE transactions on information theory, 28(2), 129–137.

Mante, V., Sussillo, D., Shenoy, K. V., & Newsome, W. T. (2013). Context-dependent computation by recurrent dynamics in prefrontal cortex. nature, 503(7474), 78–84.

Masse, N. Y., Grant, G. D., & Freedman, D. J. (2018). Alleviating catastrophic forgetting using context-dependent gating and synaptic stabilization. Proceedings of the National Academy of Sciences, 115(44), E10467–E10475.

McAdams, C. J., & Maunsell, J. H. (1999, Aug). Effects of attention on the reliability of individual neurons in monkey visual cortex. Neuron, 23(4), 765–773.

Menon, V., & D’Esposito, M. (2022). The role of pfc networks in cognitive control and executive function. Neuropsychopharmacology, 47(1), 90–103.

Miller, E. K., & Cohen, J. D. e. a. (2001). An integrative theory of prefrontal cortex function. Annual review of neuroscience, 24(1), 167–202.

Monsell, S. (2003). Task switching. Trends in cognitive sciences, 7(3), 134–140.

Motter, B. C. (1993, Sep). Focal attention produces spatially selective processing in visual cortical areas V1, V2, and V4 in the presence of competing stimuli. J Neurophysiol, 70(3), 909–919.

Musslick, S., Saxe, A., Özcimder, K., Dey, B., Henselman, G., & Cohen, J. D. (2017). Multitasking capability versus learning efficiency in neural network architectures..

Papyan, V., Han, X. Y., & Donoho, D. L. (2020). Prevalence of neural collapse during the terminal phase of deep learning training. Proceedings of the National Academy of Sciences, 117(40), 24652–24663.

Paszke, A., Gross, S., Massa, F., Lerer, A., Bradbury, J., Chanan, G., … others (2019). Pytorch: An imperative style, high-performance deep learning library. Advances in neural information processing systems, 32.

Pedregosa, F., Varoquaux, G., Gramfort, A., Michel, V., Thirion, B., Grisel, O., … Duchesnay, E. (2011). Scikit-learn: Machine learning in Python. Journal of Machine Learning Research, 12, 2825–2830.

Radford, A., Kim, J. W., Hallacy, C., Ramesh, A., Goh, G., Agarwal, S., … others (2021). Learning transferable visual models from natural language supervision. In International conference on machine learning (pp. 8748–8763).

Ravi, S., Musslick, S., Hamin, M., Willke, T. L., & Cohen, J. D. (2020). Navigating the trade-off between multi-task learning and learning to multitask in deep neural networks. arXiv preprint 2007.10527.

Reverberi, C., Görgen, K., & Haynes, J.-D. (2012). Compositionality of rule representations in human prefrontal cortex. Cerebral cortex, 22(6), 1237–1246.

Richards, B. A., Lillicrap, T. P., Beaudoin, P., Bengio, Y., Bogacz, R., Christensen, A., … others (2019). A deep learning framework for neuroscience. Nature neuroscience, 22(11), 1761–1770.

Rigotti, M., Barak, O., Warden, M. R., Wang, X.-J., Daw, N. D., Miller, E. K., & Fusi, S. (2013). The importance of mixed selectivity in complex cognitive tasks. Nature, 497(7451), 585–590.

Rosenbaum, C., Klinger, T., & Riemer, M. (2017). Routing networks: Adaptive selection of non-linear functions for multi-task learning. arXiv preprint 1711.01239.

Rougier, N. P., & O’Reilly, R. C. (2002). Learning representations in a gated prefrontal cortex model of dynamic task switching. Cognitive Science, 26(4), 503–520.

Rousselet, G. A., Fabre-Thorpe, M., & Thorpe, S. J. (2002). Parallel processing in high-level categorization of natural images. Nature neuroscience, 5(7), 629–630.

Ruder, S. (2017). An overview of multi-task learning in deep neural networks. arXiv preprint 1706.05098.

Rumelhart, D. E., Hinton, G. E., & Williams, R. J. (1986). Learning representations by back-propagating errors. nature, 323(6088), 533–536.

Samek, W., & Müller, K.-R. (2019). Towards explainable artificial intelligence. Explainable AI: interpreting, explaining and visualizing deep learning, 5–22.

Saxena, S., & Cunningham, J. P. (2019). Towards the neural population doctrine. Current opinion in neurobiology, 55, 103–111.

Serra, J., Suris, D., Miron, M., & Karatzoglou, A. (2018). Overcoming catastrophic forgetting with hard attention to the task. In International conference on machine learning (pp. 4548–4557).

Sigman, M., & Dehaene, S. (2008). Brain mechanisms of serial and parallel processing during dual-task performance. Journal of Neuroscience, 28(30), 7585–7598.

Sun, G., Probst, T., Paudel, D. P., Popović, N., Kanakis, M., Patel, J., … Van Gool, L. (2021). Task switching network for multi-task learning. In Proceedings of the ieee/cvf international conference on computer vision (pp. 8291–8300).

Vaidya, A. R., Jones, H. M., Castillo, J., & Badre, D. (2021). Neural representation of abstract task structure during generalization. ELife, 10, e63226.

Verbeke, P., & Verguts, T. (2022). Using top-down modulation to optimally balance shared versus separated task representations. Neural networks, 146, 256–271.

Vilarroya, O. (2017). Neural representation. a survey-based analysis of the notion. Frontiers in psychology, 8, 1458.

Wortsman, M., Ramanujan, V., Liu, R., Kembhavi, A., Rastegari, M., Yosinski, J., & Farhadi, A. (2020). Supermasks in superposition. Advances in Neural Information Processing Systems, 33, 15173–15184.

Yang, G. R., Joglekar, M. R., Song, H. F., Newsome, W. T., & Wang, X.-J. (2019). Task representations in neural networks trained to perform many cognitive tasks. Nature neuroscience, 22(2), 297–306.

Yuste, R. (2015). From the neuron doctrine to neural networks. Nature reviews neuroscience, 16(8), 487–497.

